# Auxiliary data, quality assurance and quality control for wearable light loggers and optical radiation dosimeters

**DOI:** 10.1101/2025.09.11.675633

**Authors:** Johannes Zauner, Oliver Stefani, Gianfranco Bocanegra, Carolina Guidolin, Björn Schrader, Ljiljana Udovicic, Manuel Spitschan

## Abstract

Wearable light loggers and optical radiation dosimeters are increasingly used in chronobiology and circadian health research, yet their data often lack contextual information (e.g., sleep, activity, environmental conditions) and may be compromised by non-wear periods, compliance issues, or technical faults. To address these limitations, we conducted interviews (n=21) and a survey (n=16) with domain experts to distill and iteratively develop auxiliary data and quality-control strategies aimed at improving the accuracy and interpretability of wearable light measurements. From this process, we established a six-domain auxiliary data framework encompassing wear/non-wear logging, sleep monitoring, light-source context, participant behaviour, user experience, and environmental light levels. Survey responses showed strong consensus on the value of auxiliary information (mean importance 4.0/5), with sleep and wear-time tracking rated as the most essential additions. To support practical adoption, we provide implementation tools, including extensions to the open-source R package LightLogR, enabling streamlined integration of wearable and auxiliary data as well as systematic quality assurance and control. Experts agreed that combining contextual records with rigorous QA/QC procedures substantially improves the reliability of field-collected light-exposure data. These recommendations and tools aim to help researchers in chronobiology, wearable sensing, and health sciences maximise data quality and enhance interpretation in real-world light-exposure studies.

## Introduction

Light affects humans beyond the visual aspect through autonomous pathways originating in the eye ^1,2^. These so-called non-visual or non-image-forming (NIF) effects of ocular light play a significant role in regulating circadian rhythms, such as the 24-hour sleep-wake cycle, and directly or indirectly affect neurological, endocrine, metabolic, and immune systems ^3^. Ideally, ocular light exposure should follow the natural light-dark cycle and its spectral and spatial properties. However, people in modern societies are often exposed to too little light during the day due to predominantly indoor activities and insufficient outdoor time. They are also exposed to too much light at night, for example, during night shift work. ^4–7^ To best support physiology, sleep, and wakefulness in healthy adults, recommendations for daytime, evening, and nighttime indoor light exposure have been published ^8^. However, little is known about human light exposure under naturalistic conditions, and a growing but fragmented landscape of researchers is taking on the task to fill this gap ^9^.

In order to assess the negative impact of modern living conditions on well-being, performance, and health, studies are required that measure personal light exposure and link these measurements to health-related outcomes. By now, studies have shown relationships of personal light exposure patterns with personal well-being and pathological outcomes, such as obesity and type II diabetes, major depression, anxiety, seasonal depressive disorder, and other pathologies related to metabolic and mental health ^10–19^.

The effects of light on human health and well-being have historically been studied in laboratory settings, examining mechanisms and dose- or phase-response relationships ^20^, and from studies over the past two decades, many principled relationships and mechanisms have been established in connecting ocular light exposure to health and health-related outcomes ^8,20^. While these highly controlled studies remain the gold standard for establishing causal and mechanistic relationships, the relevance and magnitude of real-life effects are best studied through personal light exposure data in field studies, measured through wearable devices ^21,22^. For example, laboratory studies revealing mechanistic effects of evening display light on human physiology (e.g., melanopic irradiance dependent effects on sleep latency, melatonin suppression, and alertness)^23,24^ could be modulated by prior light history, arguing for careful assessment and reporting of pre-exposure conditions ^25^. By contrast, real-world outcomes are shaped by heterogeneous daylight exposure, content/arousal, etc, which can attenuate or outweigh isolated light-mechanism effects. Recent field-oriented reviews therefore down-weight pure “brightness/arousal” accounts and emphasize usage patterns when explaining sleep impacts ^26^.

When combined with physiological measurements, wearable data contribute to understanding how laboratory findings can be translated and applied to everyday environments ^4,11,15^.

An increasing number of studies are focusing on or utilising wearable light loggers (**Figure 1A**), ranging from small but comprehensively recorded samples ^4,5,10,16,27–32^ to hundreds or even tens of thousands of participants in large cohort studies ^11,13,14,33–36^. This development is mirrored by the continued development of new devices that improve and iterate on form factors, measurement fidelity, miniaturisation, battery life, and storage capacity (**Figure 1B**)^37^. Wearable designs have also found their way into laboratory studies. As prior light history may influence the outcome of laboratory experiments investigating non-visual light effects, sleep, or chronobiology in general, it is recommended to monitor and report light exposure using wearable light loggers even before the actual experiment ^21,38^. The studies employ wearable devices to measure or estimate the ocular exposure to light and other spectral ranges of optical radiation. The devices are most often attached to a participant’s wrist, at chest level, or, most optimally, positioned at eye level ^21,39,40^. The output of these devices is commonly a time series of photopic illuminance, but depending on the device, other quantities are stored as well ^41^. These can include alpha-opic quantities like melanopic EDI, activity, temperature, direct sensor outputs, or even (reconstructed) spectral power distributions ^37,42^.

**Figure 1.**
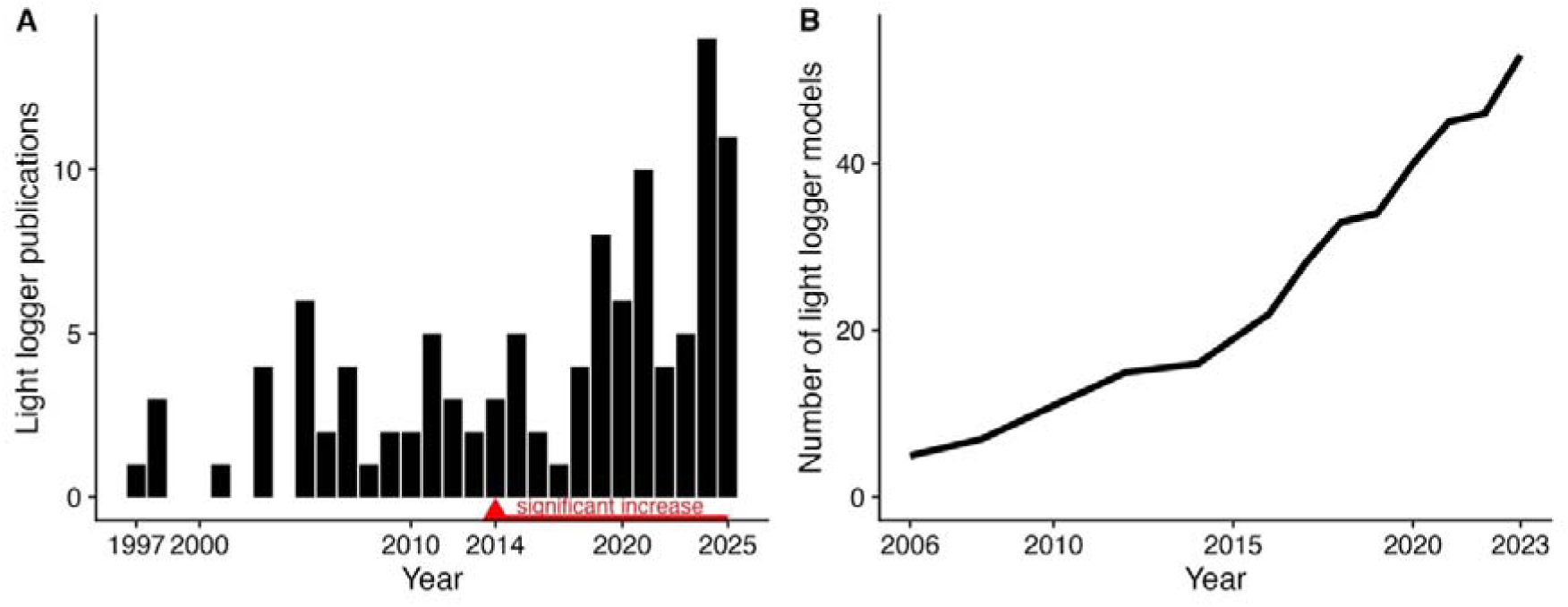
<u>Increasing relevance of light loggers in scientific publications. Data, *PubMed* search terms, and analysis script are provided in</u> **Supplementary information S2**. <u>(A) Black bars indicate the number of publications per year with reference to light loggers. The numbers are from a *PubMed* search. Only original research on humans was considered. A generalised additive model (GAM) of the publications per year reveals a significant increase in publications starting in 2014 and continuing until 2025 (indicated in red). (B) Cumulative number of light logger models on the market / used in research. Reconstructed with permission from van Duijnhoven, et al. ^37^</u>.

Furthermore, light recordings from wearables on their own – even if they are of high fidelity from a measurement perspective – often lack essential information beyond light about the environment and the participant, e.g., non-wear time, physical activity, or climate-based information ^37^. This means that additional measures are regularly required to contextualise and categorise the light and optical radiation data, which can be used either as covariates in the analysis or as quality indicators in data preprocessing, such as to detect non-wear ^43^. Beyond collecting new data, essential steps must be taken to ensure data quality throughout a study. Data from wearables can easily be compromised, for example, by clothing covering the device or if a device’s internal clock is not properly synchronised with local time. These and other confounding aspects can be mitigated through various strategies used during the study design, data collection, and data analysis, which significantly affect the utility and actual accuracy of the data ^31,41,43^. Strategies include factors such as how participants are briefed and motivated, the device placement, and the choice of analysis software.

In this study, we identified factors that can influence data quality from wearable devices used to measure light exposure and optical radiation. We also gathered and developed strategies to improve the utility of the collected data. The study is part of the MeLiDos project (Metrology for wearable light loggers and optical radiation dosimeters), dedicated to providing tools, standards, best practice guidelines, and FAIR (findable, accessible, interoperable, and reusable) data for the collection and analysis of wearables in research studies ^9^. In interviews performed by a MeLiDos subtask group with researchers familiar with the use of wearable light loggers, the experts expressed doubts about wearer compliance with the devices and the reliability of the recorded data ^44^.

This article focuses on the most important measures that can help to augment wearable data, beyond obvious technical aspects (e.g., measurement accuracy). We prioritized these because technical aspects are thoroughly treated in prior reviews ^42^, while these complementary factors emerged as under-addressed yet high-impact areas for improving data quality. The measures are divided into auxiliary data (extending the data beyond the wearable itself), quality assurance measures (increasing the likelihood of good compliance), and quality control (reducing the number of faulty data points). The auxiliary measures were surveyed and refined with researchers from across the field. The mitigating strategies were derived from discussions of common issues experienced by consortium members of the MeLiDos project and expert researcher interviews ^44^. This publication summarises the outcomes and provides guidance for auxiliary data and mitigation strategies to maximise the utility when using wearable devices in a research study.

## Results

### Auxiliary data

#### Definition

Auxiliary data in the context of this study is defined as time-dependent (i.e. time-stamped) data relevant to the analysis of light logging data but not automatically collected through wearable light logger devices, as visualised in **Figure 2**.

**Figure 2.**
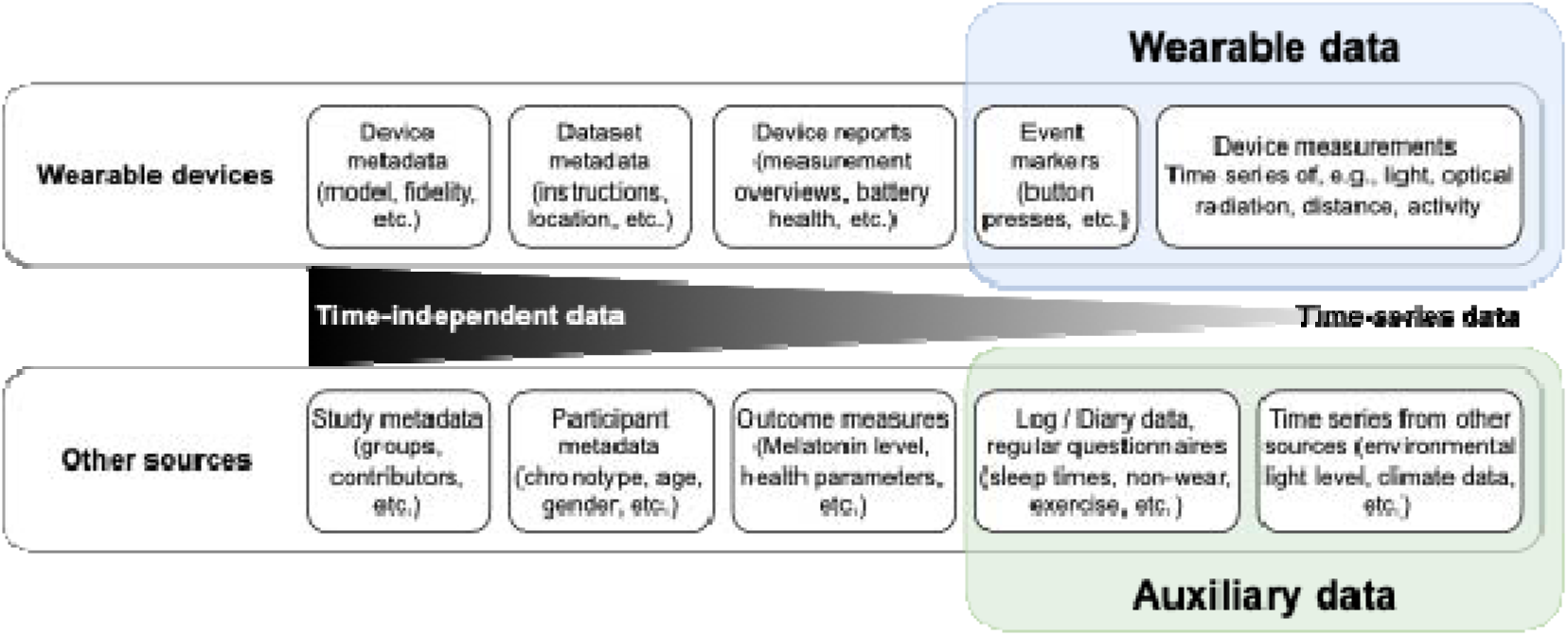
Schematic of auxiliary data in the context of other study data. Data collected in any field study using wearable devices sit on a spectrum of time-independent data (i.e., are valid independent of the time these data were recorded, like participant height or eye colour), and time-dependent data (i.e., which are valid/relevant only for part of the time, like an illuminance measurement). Some of these data can be considered metadata of the study, while others are part of the main data collection effort, e.g., as dependent variables or confounder controls. A host of secondary data that should be considered are auxiliary data, which sit on the same end of the time-dependency spectrum as data from wearables and give context to the wearable data. These data can help to filter and validate wearable data during preprocessing and allow for additional covariates during analysis.

Two simple examples are sleep-wake logs and non-wear diaries. **Figure 3** shows a visualised example of how auxiliary data extends the utility of personal light exposure data from a wearable.

**Figure 3.**
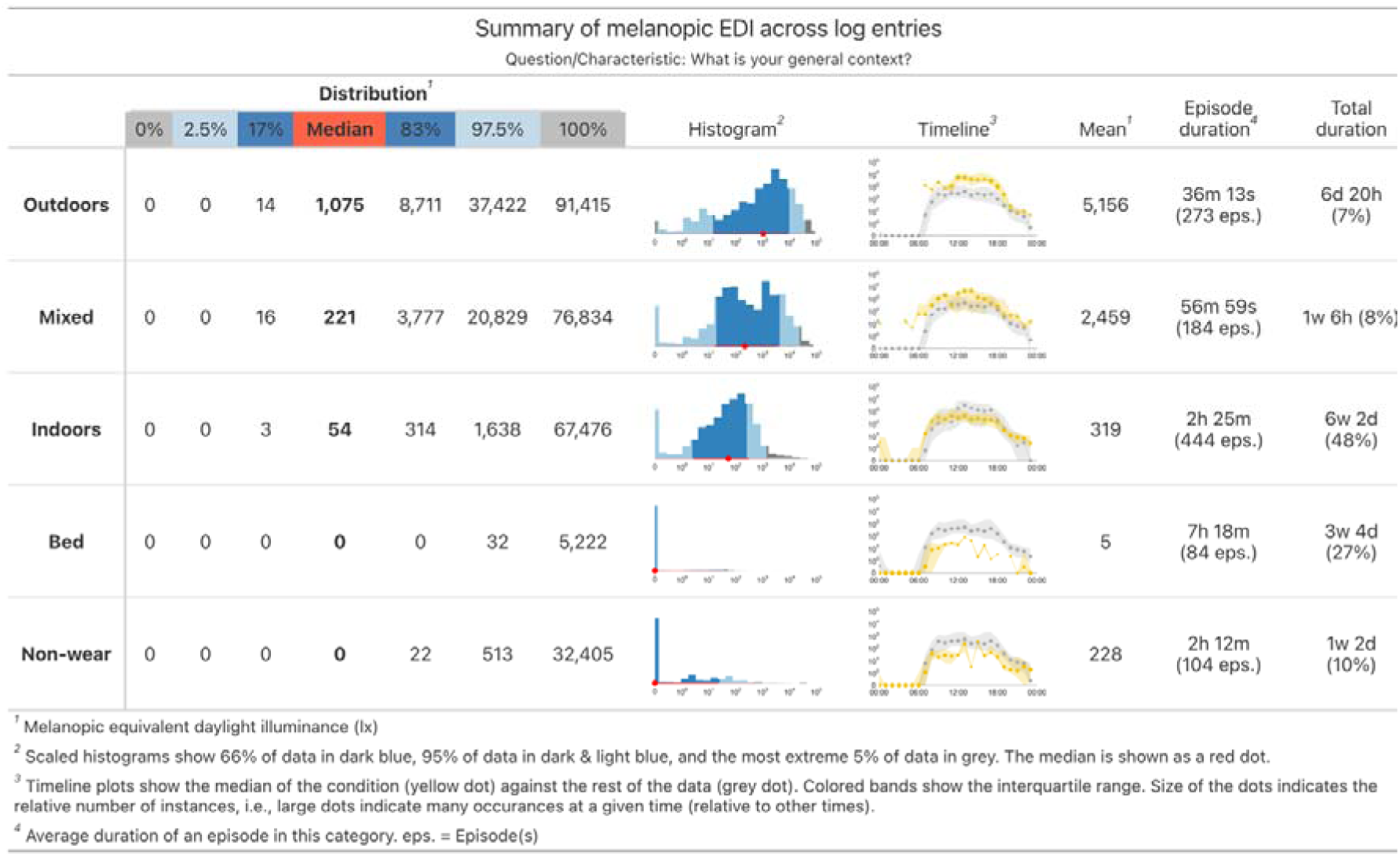
Example of the effect of auxiliary data to contextualise personal light exposure. The figure-table shows distributional characteristics of the melanopic EDI time series, a histogram, timeline plot, the mean, the mean duration of the states and number of episodes, as well as total duration for five conditions (Outdoors, Mixed indoor-outdoor, Indoors, Bedtime, and non-wear time. The effect of the auxiliary data is especially visible in the timeline plots, where the light exposure during specific states, collected through methods in the auxiliary data framework, vary strongly from the rest of the data (yellow vs. grey).

**Scope and outcome format**: All instruments in this section consist of questionnaire items and/or a set of instructions for participants. Applying these in a study result in a list of state changes connected to timestamps and participants. A simple example is shown in Table 1. The tracked states are:

- wear / non-wear
- sleep / wake
- light environment
- activity

**Table 1.**
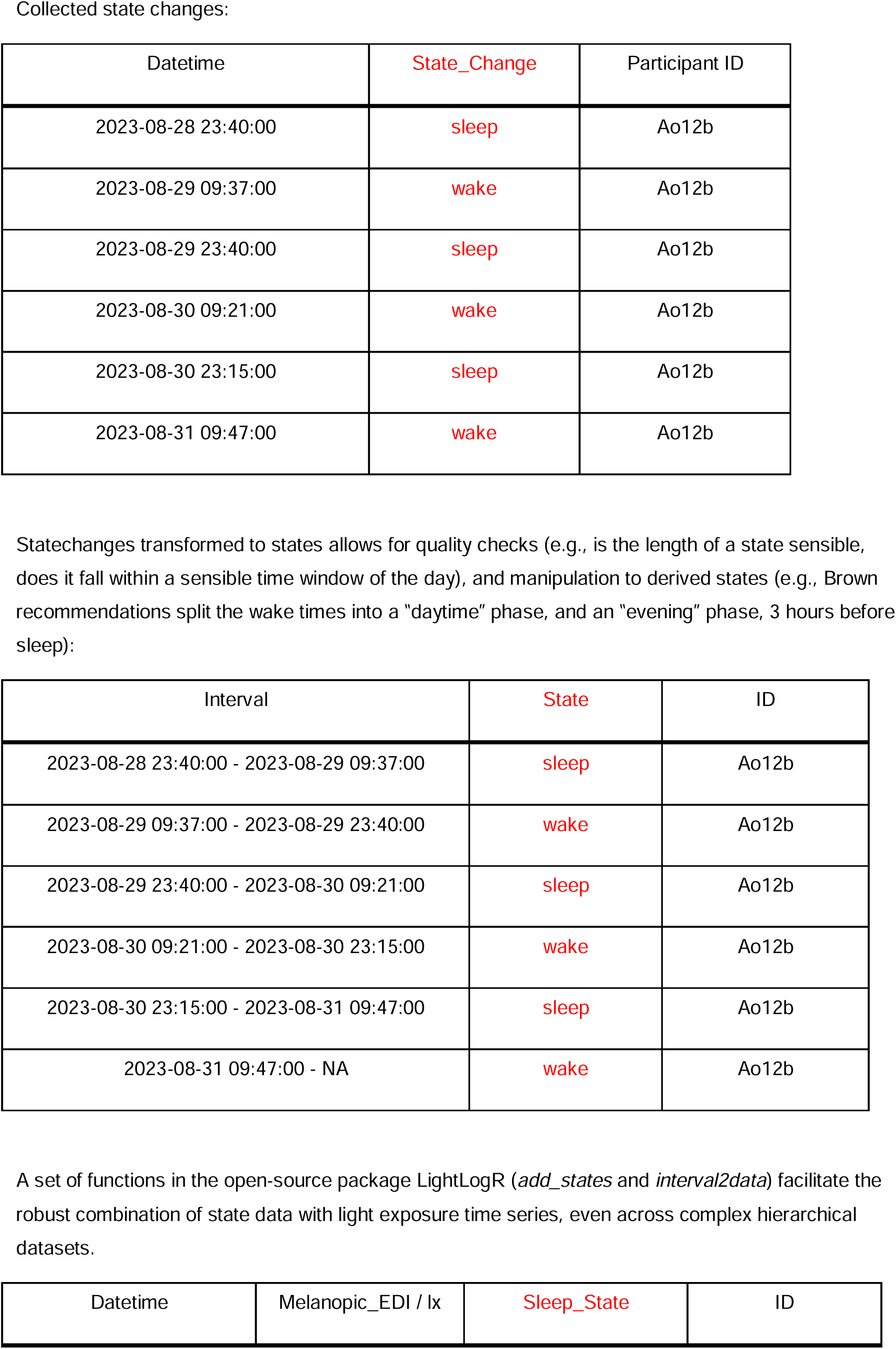
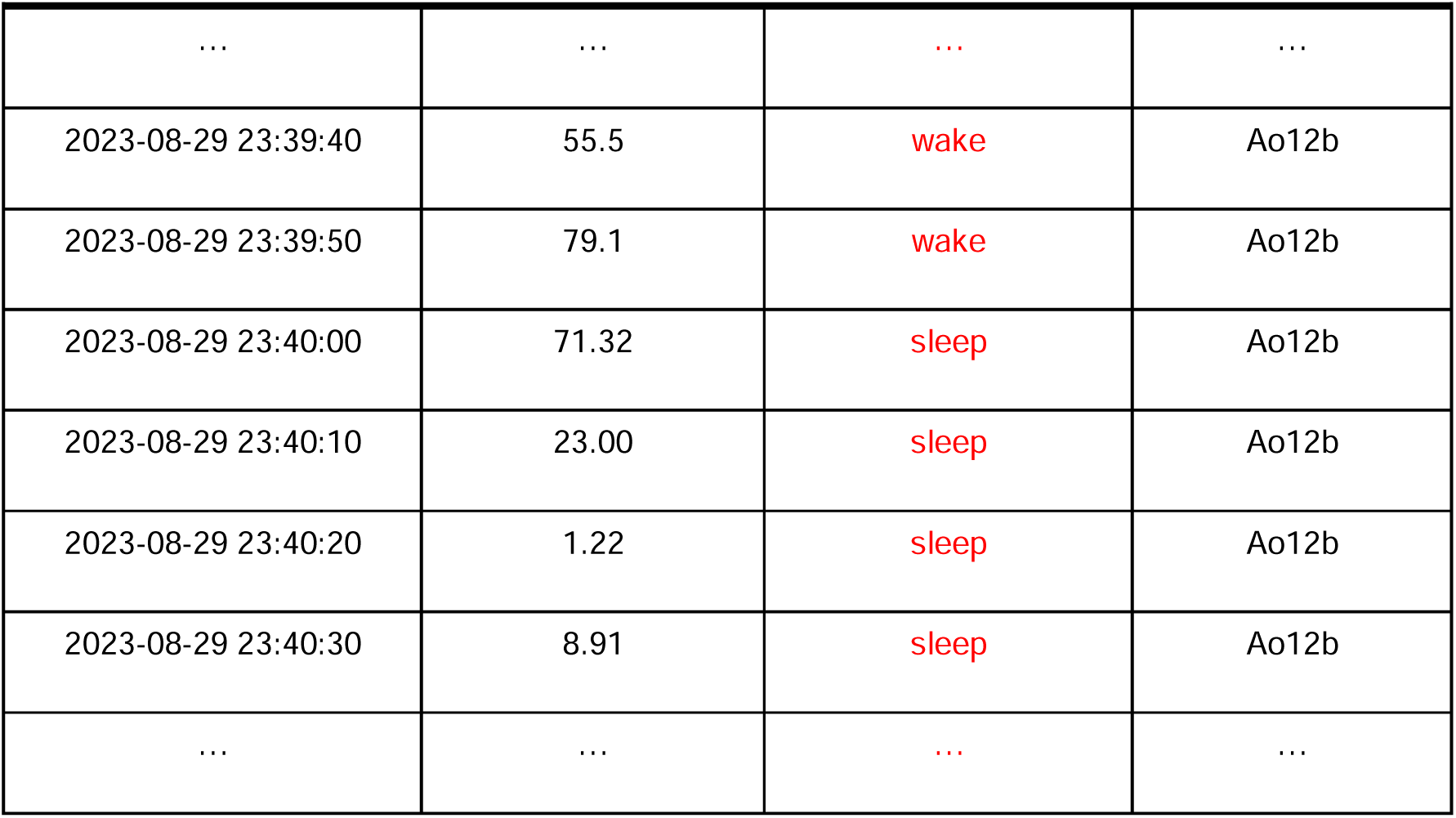
Example table for collecting state changes in sleep/wake for one participant and considerations on how these relate to actual measurements of light exposure.

Some states may be tracked already through the wearable light logger or other devices participants use. These would not be considered auxiliary data as per the definition above. In that case, the researcher must decide whether the redundant tracking through the auxiliary data strategy adds enough benefit during analysis for a particular state.

The tracked state changes can be converted to timespans of states, which relate to measurements of light and augment their usage during analysis. The MeLiDos funded software package LightLogR ^41^ features functions specifically to importing, handling, and merging auxiliary data to data from wearable light loggers in an easy and robust fashion. This includes an optional setting for upper thresholds to the length a state can sensibly persist, like 24 hours for a single wake period. The package was recently updated to version 0.10.0 (“High noon”) with functionality specifically targeted at auxiliary data and states.

### Wear log

**Objective**: Track wear and non-wear times throughout the day. To gain significant confidence, data from multiple measures should be combined.

#### Suggested measures

- A digital logbook with several options, as laid out in the procedure
- A black, light-sealed bag or container to put the wearable light logger into during non-wear time while awake (note: auxiliary data would be times of zero-lux measurements during wake times) unless light measurement outside wear time is needed for context information
- An event trigger/button on the wearable light loggers

**Possible procedure:** Throughout the day, participants are instructed to report their wear and non-wear time in a digital logbook, e.g. through a smartphone. Specifically, they have five choices of wear log entry:

- 1 = “Taking the light glasses off”
- 2 = “Putting the light glasses on”
- 3 = “Taking the light glasses off before sleep and placing them on a nightstand or flat surface”
- 4 = “Leaving <STUDY location > and its surroundings (e.g. 60 km radius)”
- 5 = “Re-entering < study location > and its surroundings (e.g. 60 km radius)”.

In case of 1, they should put their device into the black bag or container provided by the investigator. If an event button on the light logger exists and is not utilised in another way, for options 1 to 3, participants may also press the button to signal an event occurring, and in the case of 1, they are asked to confirm whether they place the light glasses in a black bag provided to them and if they are in movement.

Options 4 and 5 are introduced to control for potential differences in personal light exposure due to environmental availability rather than behaviour. They might be dropped if they are too specific for a study. For all five wear log entry choices, participants must state whether they are logging a present or a past event. That way, a lapse in procedure can be corrected by the participant, and the investigator can decide whether and under which circumstances to accept past events.

To increase compliance, a researcher might consider allowing certain non-wear periods without requiring logging (e.g., one minute).

**Utility**: Tracking non-wear time allows for the invalidation of light exposure data time periods that do not capture personal light exposure.

**Further information**: The data repository ^45^ contains one possible implementation for the wear log as it was used for a data collection effort in Tübingen, Germany.

Potentially, a sixth option could be added: “Putting the light glasses on upon waking up”.

### Morning sleep log

**Objective**: Collect sleep and wake times for each day, as well as some additional sleep-related information

#### Suggested measures

- A digital logbook with several options, as laid out in the procedure

**Possible procedure:** Every morning after waking up, participants fill in the core Consensus Sleep Diary ^46^ consisting of 9 items to assess their sleep timing, sleep duration during the night and their subjective sleep quality. This last item is scored on a five-point scale (1 = “Very poor” to 5 = “Very good”).

A researcher might decide to only include sleep/wake timing to minimise the burden on participants. This already allows the utility as described below.

**Utility**: Collecting sleep and wake times allows for metric calculation based on sleep-wake episodes instead of the 24-hour day, which can be more meaningful for certain metrics. It further allows for an assessment of how close a participant’s light exposure is to the Brown, et al. ^8^ recommendations based on daytime, pre-sleep, and sleep times.

**Further information**: The data repository ^45^ contains one possible implementation of the digital logbook, which was used for a data collection effort in Tübingen, Germany.

The morning sleep log will have to be adjusted (and renamed) for shift workers, where participants can be queried after each sleep period. Similarly, for certain research questions, capturing naptimes might be important. In this case, the log could be adjusted.

Some wearable devices can be worn during sleep, e.g. by using wristbands. These devices may track sleep-wake times automatically through actigraphy. However, they increase the error relating to the measure of interest, i.e., corneal exposure.

### Light exposure diary

**Objective**: Collect types of light environments that a participant occupied throughout the day.

#### Suggested measures

- An analogue form as laid out in the procedure
- A digital logbook to confirm compliance as laid out in the procedure
- Alternatively, a digital log might be used throughout the day for an ecological momentary assessment. However, this can affect the lighting environment in a significant way during night times (i.e, in a dark environment, there would always be partly “smartphone” or “laptop” light)

**Possible procedure**: Every evening, participants must fill in a modified version of the Harvard Light Exposure Assessment (H-LEA ^47^). This is referred to as “mH-LEA” and is done on paper using a form provided by the experimenter during the in-person visit at the start of the study. Participants are asked to report, for each hour of the day, the main light source they are exposed to. The main light source is described as “the main light source in their environment”. They can choose between 8 light categories:

- L = “Electric light source indoors (e.g.: lamps such as LEDs)”
- S = “Electric light source outdoors (e.g.: street lights)”
- I = “Daylight indoors (through windows)”
- O = “Daylight outdoors (including being in the shade)”
- E = “Emissive displays (e.g.: smartphone, laptop etc.)”
- D = “Darkness (outdoors and/or indoors)”
- W = “Light entering from outside during sleep (e.g., daylight, streetlights)”).

If they believe they are exposed to a significant combination of lights within the same hour, they can choose from the following combinations:

- “L+I”
- “L+E”
- “I+E”
- “S+O”
- “D+W”

To ensure that participants complete this task, they send a picture of the completed form every night and upload it to a shared folder (separate for each participant) where the experimenter can check compliance. Furthermore, they are asked to rate their confidence in their answers (“How sure are you about the light exposure and

activity categories you chose?”), where they can answer using a 5-point scale ranging from 1 = “Not confident at all” to 5 = “Completely confident”.

Alternatively, a digital form could be used to capture the mH-LEA directly. However, as mentioned above, this can affect the lighting environment in a significant way during night times (i.e, in a dark environment, there would always be partly “smartphone” or “laptop” light)

**Utility**: Allows for connecting subjective assessments of environmental light conditions to objective light exposure.

**Further information**: *Supplementary information 1 (C)* shows one possible implementation for the digital logbook and the analogue form, as it was used for a data collection effort in Tübingen, Germany.

The diary might be split up into two sections, each covering 12 hours instead of 24. This can reduce recollection errors by the participants.

### Behaviour/Exercise diary

**Objective**: Collect types of exercise throughout the day.

#### Suggested measures

- An analogue form as laid out in the procedure
- A digital logbook to confirm compliance as laid out in the procedure
- Alternatively, a digital log might be used throughout the day for an ecological momentary assessment.

**Possible procedure**: Every evening, participants are required to complete an experience diary form. This is done on paper using a form provided by the experimenter during the in-person visit at the start of the study. Participants are asked to report, for each hour of the day, the activity they performed in that hour. With regards to their activity, they could choose between 8 categories

- 1 = “Sleeping in bed”
- 2 = “Awake at home”
- 3 = “On the road with public transport/car”
- 4 = “On the road with bike/on foot”
- 5 = “Working in the office/from home”
- 6 = “Working outdoors (including lunch break outdoors)
- 7 = “Free time outdoors (e.g. garden/park etc.)
- 8 = “Other: please specify (e.g. sport)”

To ensure that participants complete this task, they send a picture of the completed form every night and upload it to a shared folder (separate for each participant) where the experimenter can check compliance. Furthermore, they are asked to rate their confidence in their answers (“How sure are you about the light exposure and activity categories you chose?”), where they can answer using a 5-point scale ranging from 1 = “Not confident at all” to 5 = “Completely confident”.

Alternatively, a digital form could be used to directly capture the exercise diary.

**Utility**: Allows the connection of different types of activity to objective light exposure.

**Further information**: The data repository ^45^ contains one possible implementation for the digital logbook and the analogue form, as it was used for a data collection effort in Tübingen, Germany.

### Experience log

**Objective**: Collect positive and negative experiences associated with the wearable

#### Suggested measures

- A digital logbook to confirm compliance as laid out in the procedure

**Possible procedure:** At any time, participants can note down positive and negative experiences associated with using the wearable in a digital logbook. The questions cover a wide range from what kind of experience, where it occurred and who was involved, to whether the experience resulted in any behavioural change.

**Utility:** It gives insights into a possible behavioural impact of the wearable on the participant, which might affect “regular, everyday” behaviour.

**Further information:** The data repository ^45^ contains one possible implementation for the digital logbook as it was used for a data collection effort in Tübingen, Germany.

### Environmental light levels

**Objective:** Collect objective comparative light levels that reflect the natural lighting conditions in the study location of a participant.

#### Suggested measures

- A calibrated light logger of the same type and model as used by the participant, mounted in a fixed, unobstructed position, such as a high-up rooftop. This is the measure as laid out in the procedure.
- Estimates of ground-level illuminance based on, e.g., satellite data or weather and time data. Unobstructed outdoor light levels can be derived from data provided by meteorological institutes, which typically measure global irradiance and allow for relatively straightforward conversion into illuminance values.

**Possible procedure:** The rooftop-mounted light logger collects light data during the same period as a participant. If the sensor is not rated for outdoor use, the sensor should have a clear cover, e.g., a plastic or glass dome (see **Figure 4**). We recommend a horizontal mount because the sensor can only generally reflect the environmental exposure potential.

**Figure 4.**
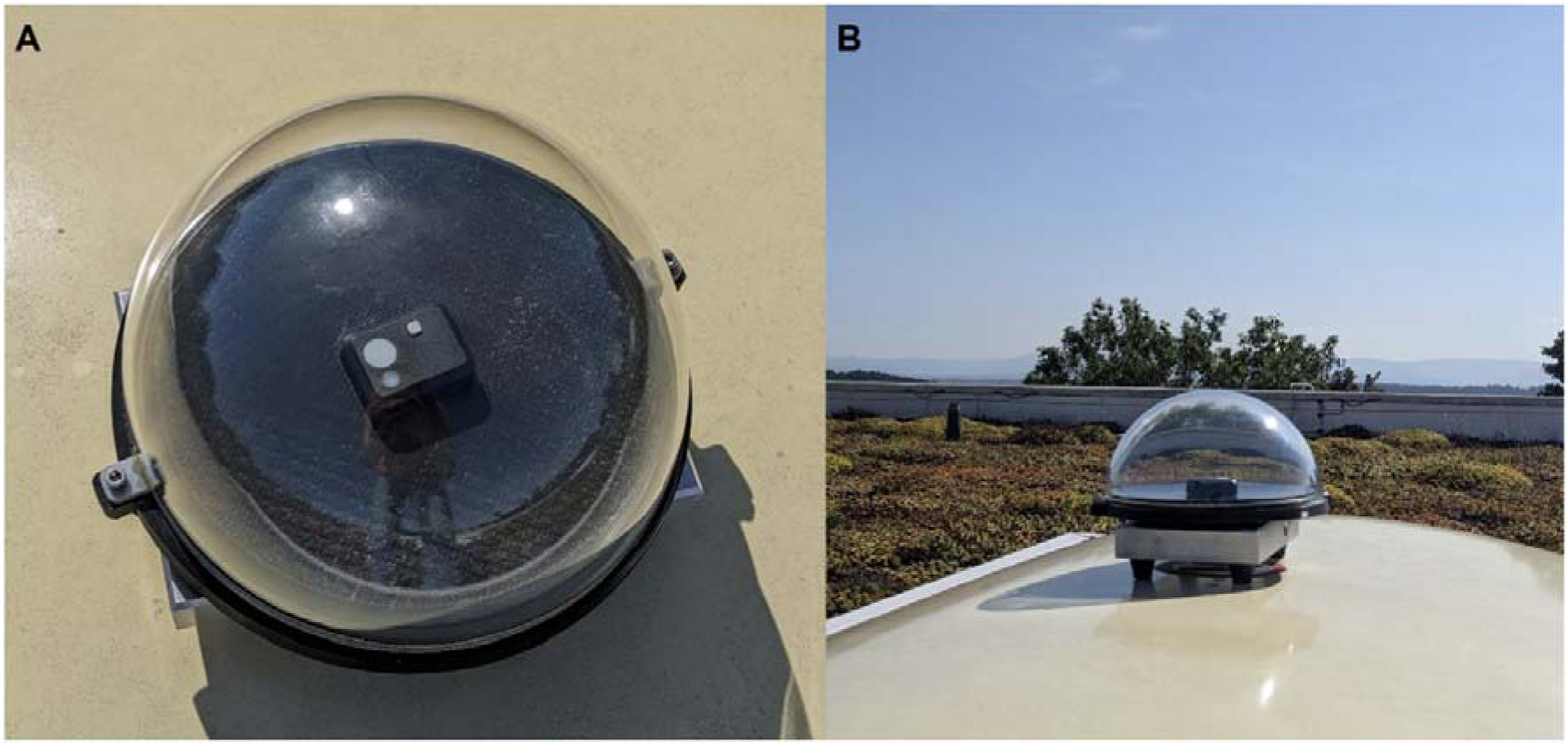
Rooftop set-up for environmental light logging. A: ActLumus light logger (Condor Instruments) horizontally placed in water-resistant set-up. B: Position of the set-up on the rooftop during measurements.

**Utility:** Gives insights into the relative light exposure a participant got compared to what would be available under natural conditions in this area. While it cannot precisely re-cap the environmental light levels at a participant’s exact position, it can do so generally.

**Further information**: *LightLogR* ^41^ has functions to connect and compare reference light levels to personal light exposure data from participants, regardless of whether they were collected with the same measurement epoch or equal time points.

For additional information, weather and ambient temperature data can provide vital context to certain behaviours affecting personal light exposure.

### Auxiliary data survey

Sixteen researchers participated in a survey collecting feedback on auxiliary data across 28 questions. They had the option to skip questions. Thus, the number of respondents (N) is provided for each question.

**Q1, agreement with the definition of auxiliary data (N=15):** All participants agreed with the definition of auxiliary data as described, with one participant adding that the utility of the experience questionnaire is not clear with regards to how it relates to the main light logger data.

It is of note that the definition of auxiliary data was adjusted slightly to cover data that are not recorded **automatically** by a device. Otherwise, event triggers or zero-lux readings enforced through the black bag would not be covered.

**Q2, previous experience with auxiliary data (N=16):** Thirteen participants collected auxiliary data before, and three had not. Of those that have previous experience, two collected data paper-based, six collected digitally, and five used a hybrid method of both. Participants could also provide additional information about their collection methods, and REDCap was mentioned three times, Qualtrics and MyCAP once each.

**Q3, most effective collection method (N=16):** All researchers agree that data should be collected digitally (n=8) or in a hybrid form (n=8). Votes for the hybrid form additionally mentioned that qualitative information or open-ended questions may be better collected via paper or interview, that participants should be given a choice about the modality, and that retrospective questionnaires about time points in the past day can profit from the paper form by providing a simple visualisation while filling the form in.

**Q4, the importance of auxiliary data (N=16):** Overall, the importance of auxiliary data is rated 4.0 on a 1 (isn’t required or helpful) to 5 (essential) scale. No participant indicated the lowest two scores (1 or 2), three gave it a middle rating (3), 10 participants selected the second highest option (4), and three participants believed light exposure measurements would be worthless without auxiliary data (5).

**Q5, relevance of auxiliary data (N=15):** Ranking the six categories of light exposure reveal that sleep/wake is the most relevant category for participants (average rank 5.0), followed by wear/non-wear (4.85), the light environment (4.13), a tie between behaviour/exercise and environmental light levels (3.07), and finally experience with the light logger (1.54).

**Q6, other domains of auxiliary data (N=10):** participants note other useful data would be on

- Clothing and accessories (e.g., hats), especially glasses/sunglasses
- Reasons for not wearing a device (author note: this has since been included in the wear log)
- A more detailed light environment (e.g., curtains)
- For intervention studies, feedback on expectations and acceptability of the intervention, symptoms, etc.
- Distinguishing work day and free day (author note: this could be extracted from the exercise diary)
- Weather conditions and temperature (author note: this aspect was added to the environmental light levels section)
- One researcher noted that the sleep diary might not work for shift workers (author note: this caveat was added to the morning sleep log section)

**Wear/Non-wear (Q7 N=16; Q8 N=16; Q9 N=5):** The domain is rated as important (3.6 on a scale of 1, not required or helpful, to 5, essential). Researchers believe the wear log captures that domain of auxiliary data sufficiently well (4.1 on a 1 to 5 scale, where 1 means it does not cover the domain at all, 2: not well, 3: partly, 4: sufficiently, 5: completely).

- Some researchers mentioned doubts as the log is quite detailed, thus increasing participant burden and potentially reducing compliance.
- One researcher suggested another option for “putting the light glasses on upon waking up” (author note: this was added as a potential addition to the wear log).
- Three researchers suggest that some way of automated non-wear detection would be much preferred. (author note: this has been added as an option in the wear log section)

**Sleep/Wake (Q10 N=16; Q11 N=16; Q12 N=2):** The domain is rated as important (4.0 on a scale of 1, not required or helpful, to 5, essential). Researchers believe the

morning sleep log captures that domain of auxiliary data sufficiently well (4.0 on a 1 to 5 scale, where 1 means it does not cover the domain at all, 2: not well, 3: partly, 4: sufficiently, 5: completely).

- Some researchers note that automated detection through actigraphy can be preferable.
- One researcher suggests extending it to include nap times (author note: this option has been added to the morning sleep log section).

**Light exposure diary (Q13 N=16; Q14 N=16; Q15 N=4):** The importance of the domain is rated as average (3.4 on a scale of 1, not required or helpful, to 5, essential). Researchers believe the light exposure diary captures that domain of auxiliary data sufficiently well (3.9 on a 1 to 5 scale, where 1 means it does not cover the domain at all, 2: not well, 3: partly, 4: sufficiently, 5: completely).

- Three researchers noted the high participant burden for this log,
- One researcher believed participants might not be able to assess different light contributions correctly.
- One researcher believes the log should be split in two, each covering 12 hours instead of 24 hours (author note: this option has been added to the light exposure diary section). Further, the utility might be highly dependent on the research question.

**Behaviour / Exercise log (Q16 N=16; Q17 N=16; Q18 N=3):** The importance of the domain is rated as average (3.2 on a scale of 1, not required or helpful, to 5, essential). Researchers believe the exercise log captures that domain of auxiliary data sufficiently well (4.1 on a 1 to 5 scale, where 1 means it does not cover the domain at all, 2: not well, 3: partly, 4: sufficiently, 5: completely).

- As with the previous question, there is some doubt about the utility of these data compared to the added participant burden.

**Experience log (Q19 N=16; Q20 N=16; Q21 N=3):** The importance of the domain is rated as average (3.0 on a scale of 1, not required or helpful, to 5, essential).

Researchers believe the experience log captures that domain of auxiliary data sufficiently well (3.8 on a 1 to 5 scale, where 1 means it does not cover the domain at all, 2: not well, 3: partly, 4: sufficiently, 5: completely).

- While researchers overall agree that it is a useful questionnaire, some note that it might be too complex and could be phrased more generally and with open-ended comments.
- One researcher notes that standards such as SUS and WEAR questionnaires are only partly suitable for usability testing of light loggers.
- Another researcher believes this log is only helpful in limited validation studies, and the nature of the log would make it hard to condense the data.

**Environmental light (Q22 N=15; Q23 N=15; Q24 N=2):** The domain is rated as important (3.7 on a scale of 1, not required or helpful, to 5, essential). Researchers believe the experience log captures that domain of auxiliary data sufficiently well (3.9 on a 1 to 5 scale, where 1 means it does not cover the domain at all, 2: not well, 3: partly, 4: sufficiently, 5: completely).

- Two researchers note that global illuminance levels do not precisely reflect environmental light levels,
- One researcher suggests using satellite data as an easier proxy.
- One researcher voices concerns about feasibility, while another believes it is a nice and easy addition.
- Another suggestion is to add information on the mounting (author note: satellite data and mounting information were added to the environmental light section).

**Q25 Impact on own research (N=15):** Two-thirds of the researchers (n=10) believe that the auxiliary data approach “[…] will definitely have some impact on [their] research design”. One-third (n=5) believe “it might have some impact […]”. None believe “it will have no impact […]”.

**Q26 general feedback (N=5):** Researchers note that the utility of the different domains depends heavily on the context, study, population, and other factors, and that other methods to add or get around these types of auxiliary data could be sought.

**Q27, Q28 (N=15 each):** Thirteen researchers would like to be mentioned as contributing to the auxiliary data questionnaire in reports and publications. Those who have not joined the group of authors have been added to the acknowledgements section. Two would like to remain anonymous. Fourteen researchers would be willing to be contacted again regarding auxiliary data, and one would not.

### Non-technical strategies for improving light data integrity

As outlined before, the quality of light exposure data collected by wearable light dosimeters can be significantly compromised by improper compliance and misuse, thus reducing accuracy to reflect the corneal exposure of the wearer. This highlights the importance of providing clear instructions, involving participants in training sessions and incorporating feedback mechanisms to improve compliance with study protocols. Addressing these factors is critical to obtaining accurate and reliable light exposure data, which can ultimately affect study results and outcomes. The following aspects have been collected from internal rounds of discussions and in-depth interviews with researchers who have used wearable devices in their studies. Table 2 provides a succinct overview of the topics, challenges, and possible strategies.

**Table 2.** Summary of quality assurance and quality control strategies.

| Category | Challenges | Strategies |
| --- | --- | --- |
| **Methodology (Device placement and usage)** | • Wrist placement: Less accurate due to clothing coverage and remoteness to the eye.<br>• Lanyard-worn and clip-worn devices are susceptible to occlusion by jackets.<br>• Glasses placement: More accurate but inconvenient and noticeable. | • Offer less intrusive placement options for balance between accuracy and comfort. |
| **Instructions and guidance** | • Unclear instructions.<br>• Lack of reminders.<br>• No feedback on incorrect positioning, battery status, or device inactivity. | • Provide clear instructions & simplify instructions with visual aids.<br>• Use automated reminders & provide real-time feedback. |
| **Social/Environmental factors** | • Social discomfort in public settings.<br>• Environmental conditions (e.g., weather, physical activities) affect usage. | • Use devices that are discreet.<br>• Offer guidance for handling the device in different conditions. |
| **Motivational feedback (Incentives and rewards)** | • No incentive system.<br>• Lack of feedback on performance. | • Introduce rewards (e.g., gift cards, gamification).<br>• Provide regular feedback on progress and impact. |
| **Data processing (Quality control techniques)** | • Need for clear criteria to exclude non-compliant data.<br>• Handling outlier data.<br>• Identifying periods when the device was worn improperly. | • Define exclusion criteria.<br>• Employ auxiliary data strategies<br>• Use advanced algorithms to detect improper wear. Apply robust outlier analysis. |

### Non-compliance and misuse

- Inconsistent use If participants do not wear the dosimeters consistently - whether due to forgetfulness, discomfort or lack of understanding - the data collected will be incomplete. This can lead to gaps in exposure data, making it difficult to draw accurate conclusions about light exposure patterns. Participants may forget to wear the dosimeter during critical exposure periods, resulting in data that does not accurately reflect their actual light exposure.
- Incorrect placement Improper placement of dosimeters can lead to inaccurate estimations of ocular light exposure. For instance, when worn on the wrist instead of near the eye (such as on spectacle frames), the device may primarily capture ambient rather than corneal illuminance, resulting in misleading data. Obstruction by clothing or hair can further impair the sensor’s ability to detect light accurately. This can lead to underestimation of exposure levels, particularly for light that would naturally reach the eyes.
- Equipment misuse Participants and even experimenters may not understand how to properly handle or calibrate the device. For example, if the dosimeter is switched off or not activated correctly, it won’t record any data, leading to data loss. Misleading feedback about the recording state could also lead to data loss. If experimenters do not follow the manufacturer’s instructions on how to use the device (e.g., how to adjust settings or interpret readings), they may misuse it, resulting in erroneous data collection.
- Lack of training and education Without sufficient training on the importance of proper use and compliance, participants may not realise how their actions affect data quality. This could lead to careless behaviour, such as not following instructions on how to wear the dosimeter or ignoring reminders to wear it consistently. If participants do not receive real-time feedback on whether they are using the dosimeter correctly, they may continue to make the same mistakes without realizing it. For example, a dosimeter that doesn’t provide acute feedback on incorrect positioning may lead to prolonged periods of incorrect use.
- Environmental and social factors Participants may feel self-conscious about wearing a conspicuous dosimeter in public, leading to avoidance of use in social situations. This may bias the data as the dosimeter may not be worn during important exposure periods. Participants may remove the dosimeter during certain activities (such as exercise or work), not realising that it’s important to collect data during these times. This results in incomplete datasets that do not represent true exposure levels.

### Strategies to improve the quality of data collection

- Ease of wear The placement of the dosimeter may be uncomfortable or inconvenient, especially when worn on glasses, which may cause participants to avoid using it or to wear it incorrectly. Devices that are small, lightweight and comfortable significantly increase compliance by minimising disruption to the wearer. Wearers are more likely to follow instructions if they forget they are wearing the device. Making devices waterproof or able to be worn in different positions without compromising accuracy further increases wearer comfort and compliance. Devices with long battery life reduce the burden on participants by limiting the frequency of charging, which can lead to data loss. Avoiding frequent charging or providing devices with rechargeable options that last for weeks ensures continuous data collection.
- Minimise behavioural disturbance Devices should avoid being visually intrusive or obvious, particularly around the eyes, which may cause wearers to behave unnaturally due to concerns about how they will be perceived by others. Visible or bulky devices could alter behaviour, thereby compromising the validity of the study. Discreet, near-eye level devices are preferred, but should not resemble cameras to avoid being mistaken for recording devices.
- Intrinsic and extrinsic motivation In some instances, participants may not fully grasp the importance of the study or lack intrinsic motivation to follow the instructions. If they do not understand how crucial accurate data is – and how they can contribute toward that end – they may not make the effort to use the dosimeter correctly. Therefore, educating participants about the scientific importance of the study can foster a sense of purpose, leading to higher intrinsic motivation and increased compliance. Offering financial rewards, personal feedback reports, or other incentives can provide external motivation. Wearers are more likely to follow instructions if they perceive a tangible benefit, such as receiving data on their own light exposure.
- Device misuse Participants may not have a clear understanding of the correct procedures for wearing and using the dosimeter, resulting in incorrect data collection; therefore, participants should receive thorough training on how to use the equipment, with regular check-ins or home visits by supervisors to ensure compliance. Reminders and instructions should be simple, emphasising the importance of the research and the correct use of the equipment to reinforce participant commitment. To prevent participants from changing their behaviour based on light exposure feedback, any real-time data, such as battery life or exposure levels, should only be visible to researchers. Providing this feedback only to the researcher avoids biasing the results while still ensuring the functionality of the device.

### Strategies to improve the quality of data processing: Exclusion and calibration (quality control)

- Transparency of calibration A consistent concern voiced by researchers in the interviews is the lack of transparency in how instruments are calibrated by manufacturers. Standardised calibration procedures, including inter-device comparisons and position-specific validations (e.g. wrist vs. chest), are essential to ensure reliable measurements. Improving the transparency of the calibration process will increase researchers’ confidence and improve data quality, especially in low-light or extreme conditions.
- Exclusion criteria Researchers often exclude data taken under certain conditions, such as below 20 lux, due to concerns about accuracy in low light or overly bright environments (e.g. saturated levels). Establishing clear exclusion criteria based on well-understood device limitations ensures that only reliable data is used in the analysis.
- Data cleaning and interpretation Depending on the device, participant compliance, and other factors, it is commonly necessary to invalidate some of the measurements recorded by a device. Reasons include aspects such as:

○ out of range measurements (beyond sensor saturation and at the noise level)
○ non-wear
○ device occlusion
○ recordings outside relevant time frames (trimming) Often, these invalid data are randomly distributed, but due to non-wear at times of day at which specific activities occur (e.g., showering, contact sports, swimming, …), light exposure data may be missing systematically. The following sanity checks are recommended:
○ Visualisation of raw time series for the available sensors
○ Visualisation of histograms to identify distributions of data and outliers Combining device data with participant-reported activities (e.g. diaries, apps – see the section on auxiliary data above) is a common strategy to contextualise light exposure, but such methods are considered unreliable by some due to participant forgetfulness. Automated data collection or digital wearable time recording devices coupled with motion sensors are recommended for greater accuracy. There is a need for standardisation of data cleaning methods (e.g. identification of non-wear times, processing of light data by body position). Standard protocols will ensure that data from different studies are comparable, thus improving the overall quality of research results.
- Data compression and connectivity Large datasets can lead to interruptions in downloads, creating a risk of data loss. Software developers should prioritise efficient data compression techniques without compromising quality. Reliable connectivity (Bluetooth, cable or infrared) should be emphasised to avoid transmission problems during data extraction.

## Discussion

The utility of personal light exposure data heavily depends on key decisions and strategies before, during, and after data collection. In this study, we present essential aspects, both to maximise the quality of light exposure data themselves, as well as for auxiliary data to the time series.

The auxiliary data strategy in this publication covers six domains: wear and non-wear of the devices, sleep, light sources, behaviour, experience, and environmental light levels. Sixteen experts in the field were surveyed towards this strategy. All researchers agree on the definition of auxiliary data, and most of them have previous experience collecting it. Furthermore, all surveyed participants agree that auxiliary data are important, with a third believing them “essential”. Of the different modalities of auxiliary data, sleep and wear data are considered to be the most important, but none are considered irrelevant. Importantly, the presented solutions were (on average) considered as sufficient to cover each domain, and all researchers believe the strategy will or might have some impact on their future research. This suggests a robust basis to collect auxiliary data in these domains.

Besides these encouraging results, many answers caution that some domains or the whole approach would heavily increase participant burden, a concern we share. The benefits of additional data have to be weighed against potentially reduced participant compliance, errors, or, depending on the study goal, questionable utility of a given domain. Thus, it has to be emphasised that we do not recommend every future study to include the auxiliary data strategy in full and as laid out here. Rather, we want to make researchers aware of domains of auxiliary data that have been of importance in the past and suggest a possible implementation to capture these data. During study design, each domain and measure has to be weighed against the additional burden for participants, the study population’s willingness and ability to comply, and alternative ways to collect or add the data. For example, the experience log can be a good tool to assess different wearing positions in a pilot study, but might be dropped from a larger-scale collection effort. Ideally, as devices and analysis tools become more sophisticated, the need for some of these domains will decrease, e.g., when wear times are assessed and recorded by the devices themselves. Furthermore, contextual or environmental data may become easier to collect, as interfaces to public services such as meteorological institutes become more interoperable and accessible.

Collection of non-wear data was rated as important by researchers participating in our survey. While self-reported methods (e.g., wear logs) were generally considered effective, some researchers emphasised the need for automated detection. A recent study by Guidolin et al. (2025)^43^ evaluated a low-illuminance cluster–based algorithm for detecting non-wear periods and found good agreement with self-reported wear logs (ground truth), provided that participants stored the device in a light-blocking bag during non-wear time. Importantly, their analysis showed that accounting for non-wear periods affected only half of the light exposure metrics investigated (7 of 14), and even among those, most differences were small. Significant effects were primarily observed when comparing uncorrected (raw) data to corrected datasets, rather than between correction methods. For instance, average differences between raw and wear log–corrected data for key duration metrics (TAT250, TAT1000) were minimal—approximately 2 and 1 minutes in total, respectively. These results suggest that the influence of non-wear time on derived metrics is generally modest but can become relevant in studies focusing on specific light exposure periods (e.g., evenings). In the absence of a standardised non-wear detection framework, researchers should therefore align their data collection and preprocessing approach with the specific aims and sensitivity requirements of their study. To improve the quality and usability of light exposure data, future studies should adopt participant-centred strategies, including optimised device placement, clear instructions, real-time feedback, a discreet and weather-resistant design, motivational incentives and robust data quality control, to balance measurement accuracy with compliance and comfort. There are four resources we want to highlight in that regard:

- The open-source software package LightLogR by Zauner, et al. 41 provides standardised routines to ingest data from many wearable devices, detect and handle gaps and irregularities, merge data from different sources (such as wear or sleep/wake), visualise, and perform metric calculations. Using these standard analysis pipelines not only reduces time spent preparing the data, but also reduces errors and differences when implementing routines on a by-workgroup basis. Compared to the version at the time of the referenced paper, a recent update (0.10.0 High noon) 48 has added many functions related to states, which are at the core of auxiliary data.
- A recent survey by Zauner, et al. 49 with over 150 participants from around the world explored wearer preferences regarding eight different wearing positions for wearable devices. Surveyed contexts include work, home, social, and sport, which have slight but significant effects on user preferences. This survey can be one resource to guide decisions on device type and placement based on study duration and contexts.
- A research guide by the Research data alliance (RDA) working group on optical radiation exposure and visual experience data (in active development, https://rda-wg-visualexperiencedata.github.io/ResearcherGuide/) is a rich resource for many technical and non-technical data quality aspects.
- A device specification tool, developed as another part of the MeLiDos project contains many aspects relating to data quality and wearability (https://tscnlab-wearable-devices-specification.share.connect.posit.cloud) 50. The tool combines these aspects into a specification sheet that facilitates device selection and procurement.
- The project A day in daylight 51, as part of the Daylight Awareness Week 2025 combines light exposure data from researchers around the globe (47 participants), collected on the day of the solar equinox with a rich event log (over 1700 log entries). These log entries relate to or directly encapsulate many auxiliary data concepts. While this project is not a traditional research study, and the sample is biased, the conditional light exposure tables still highlight the relevance and effect of auxiliary data. Part of that information is contained in Figure 3, but many more contexts are available in the accessible project dashboard (https://tscnlab.github.io/2025_ADayInDaylight/).

A measure not discussed in this study, but still noteworthy, is that random acts of non-compliance or misuse become less relevant for light exposure metrics as the data collection duration increases. In a bootstrap analysis of a month-long collection period per person, Biller, et al. ^6^ found that even 6 hours of missing data per day did not significantly affect light exposure metrics across the whole period, with shorter collection periods obviously more sensitive to missingness than longer ones. While missing data is not equal to measurement changes due to non-compliance or misuse, the trend towards less effect over longer periods could still hold.

There are noteworthy limitations to our study. As field studies with wearable light loggers and optical radiation dosimeters are still an emerging field of research (**Figure 1**), the suggestions and measures provided above must be considered optional rather than required. They reflect best practices available at the time of publication and need to be considered in the context of a given study protocol.

Depending on a study’s goal or scope, only some suggested measures might be useful or sensible. Participant burden is also an aspect to be considered carefully.

Further, the measures are not infallible, and, in many cases, their benefit cannot be quantified. This limitation is mostly grounded in how the measures were collected: as experience-based solutions to problems researchers encountered in their work. As such, they must be considered a byproduct of research rather than the actual topic, which would be required for a quantitative assessment of their benefit.

## Methods

### Data collection

Data, i.e. strategies and questionnaires, were collected in multiple rounds of discussion among the participating MeLiDos consortium members in 2024.

In addition, in-depth interviews were conducted with 21 researchers experienced in the use of light loggers (invited: 101; invitation accepted: 21). While the full results of these interviews will be reported separately ^44^, this paper extracts and expands upon the key findings specifically related to strategies for augmenting data quality.

Furthermore, a survey was conducted in the third quarter of 2024 for the auxiliary data. The survey was hosted on SurveyMonkey and was sent out to 106 researchers. Because the survey invitation was issued later, we added researchers to the recruitment list as they were identified or as they joined the field. Invited researchers were experts known to MeLiDos consortium members, the CIE joint technical committee JTC20, and corresponding authors from papers cited in a recent review about metrics used in wearable light logger studies by Hartmeyer and Andersen ^52^. Sixteen (n=16) researchers participated in the survey, providing additional insights and suggestions, which were consequently implemented into the results. The survey was accompanied by a preliminary version of the auxiliary data presented in this study. The 28 survey questions ranged from general questions about the nature and utility of auxiliary data to questions specifically targeted at individual aspects of auxiliary data in the preliminary document, such as sleep or non-wear. The survey results can be found in *Supplementary information S1*.

In general, both the survey and interview participants (i.e., experts and researchers) can be considered a form of convenience sample. However, given the emerging yet relatively small size of the research field (**Figure 1A**), this sample represents a substantial proportion of the known expert community (∼ 21 and 16 out of 106), considering that the invited group stemmed from a comprehensive review of the field by Hartmeyer and Andersen ^52^ with a size of only about 100 corresponding researchers.

### Data analysis

Collected data consisted of long-form interviews, questionnaires, and (mostly) free-form survey replies. Data were condensed through manual, topical analysis, and expert feedback was manually implemented.

## Supporting information

Figure 1 generation

Survey results

## Declaration statements

### Data availability

All relevant data are part of this publication, either contained in the manuscript or its supplements.

### Code availability

The auxiliary data questionnaires (PDF) and REDCap questionnaires for the *Morning sleep diary*, the *Wear log*, the *Light Exposure diary*, and the *Experience log* are available from the GitHub repository (https://github.com/tscnlab/ZaunerEtAl_npjBiolTimingSleep_2025), archived on Zenodo ^45^. The repository further contains code and data to recreate **Figure 1**.

## Acknowledgments

We would like to thank the following researchers (in alphabetical order) and three anonymous researchers that participated in the auxiliary data survey and contributed to the auxiliary data strategy as it is presented here: Tianren Chen, Débora B. Constantino, Altug Didikoglu, Fatemeh Fazlali, Gena Glickman, Steffen Hartmeyer, Rafael Lazar, Renske Lok, Elise McGlashan, Karin Smolders, and Jamie Zeitzer

The authors were financially supported by MeLiDos. The project (22NRM05 MeLiDos) has received funding from the European Partnership on Metrology, co-financed by the European Union’s Horizon Europe Research and Innovation Programme and by the Participating States.

On the use of generative AI and AI-assisted technologies in the writing process: The authors used ChatGPT during the preparation of this work. After using this tool, the authors reviewed and edited the content as needed and take full responsibility for the content of the publication.

The following use of AI by contributor roles is based on the CRediT taxonomy. Funding acquisition, project administration, resources, and supervision were deemed irrelevant in this context and thus removed:

Conceptualization: no Data curation: no Investigation: no Formal analysis: no Methodology: no Software: no Validation: no Visualization: no

Writing – original draft: abstract refinement

Writing – review & editing: improve readability and language

## Authors’ contributions

Conceptualisation: J.Z., O.S., C.G.

Data Curation: J.Z., O.S., C.G.

Formal Analysis: -

Funding Acquisition: M.S., B.S.

Investigation: J.Z., O.S., G.B.

Methodology: J.Z., O.S., C.G.

Project Administration: M.S., B.S., J.Z., O.S

Resources: -

Software: -

Supervision: M.S.

Validation: -

Visualisation: J.Z.

Writing – Original Draft Preparation: J.Z., O.S.

Writing – Review & Editing: All authors

## Supplementary information

**Supplementary information S1.** PDF document containing the (anonymous) survey results for the auxiliary data strategy. Email addresses in Q27 and Q28 have been removed for privacy reasons. Names in Q27 have been preserved, as participants opted to be named as contributors.

**Supplementary information S2.** Data and analysis script to generate Figure 1

## References

1 Berson, D. M., Dunn, F. A. & Takao, M. Phototransduction by retinal ganglion cells that set the circadian clock. Science 295, 1070–1073 (2002). 10.1126/science.1067262

2 Hattar, S., Liao, H. W., Takao, M., Berson, D. M. & Yau, K. W. Melanopsin-containing retinal ganglion cells: architecture, projections, and intrinsic photosensitivity. Science 295, 1065–1070 (2002). 10.1126/science.1069609

3 Schmidt, T. M., Chen, S. K. & Hattar, S. Intrinsically photosensitive retinal ganglion cells: many subtypes, diverse functions. Trends Neurosci 34, 572–580 (2011). 10.1016/j.tins.2011.07.001

4 Didikoglu, A., Mohammadian, N., Johnson, S., van Tongeren, M., Wright, P., Casson, A. J., Brown, T. M. & Lucas, R. J. Associations between light exposure and sleep timing and sleepiness while awake in a sample of UK adults in everyday life. Proc Natl Acad Sci U S A 120, e2301608120 (2023). 10.1073/pnas.2301608120

5 Price, L. L. A., Khazova, M. & Udovicic, L. Assessment of the Light Exposures of Shift-working Nurses in London and Dortmund in Relation to Recommendations for Sleep and Circadian Health. Ann Work Expo Health 66, 447–458 (2022). 10.1093/annweh/wxab092

6 Biller, A. M., Zauner, J., Cajochen, C., Gerle, M. A., Kalavally, V., Mohamed, A., Rottländer, L., Seah, M.-Y., Stefani, O. & Spitschan, M. Physiologically-relevant light exposure and light behaviour in Switzerland and Malaysia. Journal of Exposure Science & Environmental Epidemiology, (2025). 10.1038/s41370-025-00825-8

7 Agbeshie, G. K., Duah Junior, I. O., Andoh, A. K. A., Ampong, J., Mensah, N. A. O., Ampoma-Mensah, A. Y., Zauner, J., Spitschan, M. & Akuffo, K. O. Physiologically relevant real-world light exposure and its behavioural and environmental determinants in Kumasi, Ghana. Open Res Eur 5, 300 (2025). 10.12688/openreseurope.21304.1

8 Brown, T. M., Brainard, G. C., Cajochen, C., Czeisler, C. A., Hanifin, J. P., Lockley, S. W., Lucas, R. J., Munch, M., O’Hagan, J. B., Peirson, S. N., Price, L. L. A., Roenneberg, T., Schlangen, L. J. M., Skene, D. J., Spitschan, M. et al. Recommendations for daytime, evening, and nighttime indoor light exposure to best support physiology, sleep, and wakefulness in healthy adults. PLoS Biol 20, e3001571 (2022). 10.1371/journal.pbio.3001571

9 Spitschan, M., Zauner, J., Nilsson Tengelin, M., Bouroussis, C. A., Caspar, P. & Eloi, F. Illuminating the future of wearable light metrology: Overview of the MeLiDos Project. Measurement 235, 114909 (2024). 10.1016/j.measurement.2024.114909

10 Nowozin, C., Wahnschaffe, A., de Zeeuw, J., Papakonstantinou, A., Hadel, S., Rodenbeck, A., Bes, F. & Kunz, D. Living in Biological Darkness II: Impact of Winter Habitual Daytime Light on Night-Time Sleep. Eur J Neurosci 61, e16647 (2025). 10.1111/ejn.16647

11 Burns, A. C., Windred, D. P., Rutter, M. K., Olivier, P., Vetter, C., Saxena, R., Lane, J. M., Phillips, A. J. K. & Cain, S. W. Day and night light exposure are associated with psychiatric disorders: an objective light study in >85,000 people. Nat Ment Health 1, 853–862 (2023). 10.1038/s44220-023-00135-8

12 Gubin, D., Danilenko, K., Stefani, O., Kolomeichuk, S., Markov, A., Petrov, I., Voronin, K., Mezhakova, M., Borisenkov, M., Shigabaeva, A., Yuzhakova, N., Lobkina, S., Petrova, J., Malyugina, O., Weinert, D. et al. Light Environment of Arctic Solstices is Coupled With Melatonin Phase-Amplitude Changes and Decline of Metabolic Health. J Pineal Res 77, e70023 (2025). 10.1111/jpi.70023

13 Windred, D. P., Burns, A. C., Rutter, M. K., Ching Yeung, C. H., Lane, J. M., Xiao, Q., Saxena, R., Cain, S. W. & Phillips, A. J. K. Personal light exposure patterns and incidence of type 2 diabetes: analysis of 13 million hours of light sensor data and 670,000 person-years of prospective observation. Lancet Reg Health Eur 42, 100943 (2024). 10.1016/j.lanepe.2024.100943

14 Windred, D. P., Burns, A. C., Lane, J. M., Olivier, P., Rutter, M. K., Saxena, R., Phillips, A. J. K. & Cain, S. W. Brighter nights and darker days predict higher mortality risk: A prospective analysis of personal light exposure in >88,000 individuals. Proc Natl Acad Sci U S A 121, e2405924121 (2024). 10.1073/pnas.2405924121

15 Burns, A. C., Saxena, R., Vetter, C., Phillips, A. J. K., Lane, J. M. & Cain, S. Time spent in outdoor light is associated with mood, sleep, and circadian rhythm-related outcomes: a cross-sectional and longitudinal study in over 400,000 UK Biobank participants. J Affect Disord 295, 347–352 (2021). 10.1016/j.jad.2021.08.056

16 Gubin, D., Danilenko, K., Stefani, O., Kolomeichuk, S., Markov, A., Petrov, I., Voronin, K., Mezhakova, M., Borisenkov, M., Shigabaeva, A., Yuzhakova, N., Lobkina, S., Weinert, D. & Cornelissen, G. Blue Light and Temperature Actigraphy Measures Predicting Metabolic Health Are Linked to Melatonin Receptor Polymorphism. Biology (Basel) 13, 22 (2023). 10.3390/biology13010022

17 Leger, D., Bayon, V., Elbaz, M., Philip, P. & Choudat, D. Underexposure to light at work and its association to insomnia and sleepiness: a cross-sectional study of 13,296 workers of one transportation company. J Psychosom Res 70, 29–36 (2011). 10.1016/j.jpsychores.2010.09.006

18 Gubin, D., Julia, B., Oliver, S., Sergey, K., Liina, D., Alexander, M., Aislu, S., Germaine, C. & Weinert, D. Light exposure predicts COVID-19 negative status in young adults. Biol Rhythm Res 55, 535–546 (2024). 10.1080/09291016.2024.2427608

19 Gubin, D., Kolomeichuk, S., Danilenko, K., Stefani, O., Markov, A., Petrov, I., Voronin, K., Mezhakova, M., Borisenkov, M., Shigabaeva, A., Boldyreva, J., Petrova, J., Weinert, D. & Cornelissen, G. Light Exposure, Physical Activity, and Indigeneity Modulate Seasonal Variation in NR1D1 (REV-ERBalpha) Expression. Biology (Basel) 14, 231 (2025). 10.3390/biology14030231

20 Brown, T. M. Melanopic illuminance defines the magnitude of human circadian light responses under a wide range of conditions. J Pineal Res 69, e12655 (2020). 10.1111/jpi.12655

21 Spitschan, M., Smolders, K., Vandendriessche, B., Bent, B., Bakker, J. P., Rodriguez-Chavez, I. R. & Vetter, C. Verification, analytical validation and clinical validation (V3) of wearable dosimeters and light loggers. Digit Health 8, 20552076221144858 (2022). 10.1177/20552076221144858

22 Spitschan, M. & Joyce, D. S. Human-Centric Lighting Research and Policy in the Melanopsin Age. Policy Insights Behav Brain Sci 10, 237–246 (2023). 10.1177/23727322231196896

23 Schollhorn, I., Stefani, O., Lucas, R. J., Spitschan, M., Slawik, H. C. & Cajochen, C. Melanopic irradiance defines the impact of evening display light on sleep latency, melatonin and alertness. Commun Biol 6, 228 (2023). 10.1038/s42003-023-04598-4

24 Cajochen, C., Frey, S., Anders, D., Spati, J., Bues, M., Pross, A., Mager, R., Wirz-Justice, A. & Stefani, O. Evening exposure to a light-emitting diodes (LED)-backlit computer screen affects circadian physiology and cognitive performance. J Appl Physiol 110, 1432–1438 (2011). 10.1152/japplphysiol.00165.2011

25 Schöllhorn, I., Stefani, O., Blume, C. & Cajochen, C. Seasonal Variation in the Responsiveness of the Melanopsin System to Evening Light: Why We Should Report Season When Collecting Data in Human Sleep and Circadian Studies. Clocks Sleep 5, 651–666 (2023). 10.3390/clockssleep5040044

26 Bauducco, S., Pillion, M., Bartel, K., Reynolds, C., Kahn, M. & Gradisar, M. A bidirectional model of sleep and technology use: A theoretical review of How much, for whom, and which mechanisms. Sleep Med Rev 76, 101933 (2024). 10.1016/j.smrv.2024.101933

27 Guidolin, C., Aerts, S., Agbeshie, G. K., Akuffo, K. O., Aydin, S. N., Baeza-Moyano, D., Bolte, J., Broszio, K., Cantarero-Garcia, G., Didikoglu, A., Gonzalez-Lezcano, R. A., Joosten-Ma, H., Melero-Tur, S., Tengelin, M. N., Perez Gutierrez, M. C. et al. Protocol for a prospective, multicentre, cross-sectional cohort study to assess personal light exposure. BMC Public Health 24, 3285 (2024). 10.1186/s12889-024-20206-4

28 Estevan, I., Tassino, B., Vetter, C. & Silva, A. Bidirectional association between light exposure and sleep in adolescents. J Sleep Res 31, e13501 (2022). 10.1111/jsr.13501

29 Hand, A. J., Stone, J. E., Shen, L., Vetter, C., Cain, S. W., Bei, B. & Phillips, A. J. K. Measuring light regularity: sleep regularity is associated with regularity of light exposure in adolescents. Sleep 46, zsad001 (2023). 10.1093/sleep/zsad001

30 da Costa Lopes, L., Ribeiro da Silva Vallim, J., Tufik, S., Louzada, F. & D’Almeida, V. Associations between real-life light exposure patterns and sleep behaviour in adolescents. J Sleep Res 33, e14315 (2024). 10.1111/jsr.14315

31 Stefani, O., Marek, R., Schwarz, J., Plate, S., Zauner, J. & Schrader, B. Wearable Light Loggers in Field Conditions: Corneal Light Characteristics, User Compliance, and Acceptance. Clocks Sleep 6, 619–634 (2024).

32 Biller, A. M., Zauner, J., Cajochen, C., Gerle, M. A., Kalavally, V., Mohamed, A., Rottländer, L., Seah, M.-Y., Stefani, O. & Spitschan, M. Physiologically-relevant light exposure and light behaviour in Switzerland and Malaysia. bioRxiv, 2025.2001.2007.631760 (2025). 10.1101/2025.01.07.631760

33 Lok, R., Ancoli-Israel, S., Ensrud, K. E., Redline, S., Stone, K. L. & Zeitzer, J. M. Timing of outdoor light exposure is associated with sleep-wake consolidation in community-dwelling older men. Front Sleep 2, 1268379 (2023). 10.3389/frsle.2023.1268379

34 Li, Q., Xu, Y. X., Lu, X. Z., Shen, Y. T., Wan, Y. H., Su, P. Y., Tao, F. B., Chen, X. & Sun, Y. Impact of bedroom light exposure on glucose metabolic markers and the role of circadian-dependent meal timing: A population-based cross-sectional study. Ecotoxicol Environ Saf 290, 117589 (2024). 10.1016/j.ecoenv.2024.117589

35 Wallace, D. A. Light exposure differs by sex in the US, with females receiving less bright light. npj Biol Timing Sleep 1, 16 (2024). 10.1038/s44323-024-00016-y

36 Wallace, D. A., Redline, S., Sofer, T. & Kossowsky, J. Environmental Bright Light Exposure, Depression Symptoms, and Sleep Regularity. JAMA Netw Open 7, e2422810 (2024). 10.1001/jamanetworkopen.2024.22810

37 van Duijnhoven, J., Hartmeyer, S. L., Didikoglu, A., Stefani, O., Houser, K. W., Kalavally, V. & Spitschan, M. Measuring light exposure in daily life: A review of wearable light loggers. Build Environ 274, 112771 (2025). 10.1016/j.buildenv.2025.112771

38 Roguski, A., Needham, N., MacGillivray, T., Martinovic, J., Dhillon, B., Riha, R. L., Armstrong, L., Campbell, I. H., Ferguson, A., Hilgen, G., Lako, M., Ritter, P., Santhi, N., von Schantz, M., Spitschan, M. et al. Investigating light sensitivity in bipolar disorder (HELIOS-BD). Wellcome Open Res 9, 64 (2024). 10.12688/wellcomeopenres.20557.1

39 Hartmeyer, S. L., Webler, F. S. & Andersen, M. Towards a framework for light-dosimetry studies: Methodological considerations. Light Res Technol 55, 377–399 (2022). 10.1177/14771535221103258

40 Stampfli, J. R., Schrader, B., di Battista, C., Häfliger, R., Schälli, O., Wichmann, G., Zumbühl, C., Blattner, P., Cajochen, C., Lazar, R. & Spitschan, M. The Light-Dosimeter: A new device to help advance research on the non-visual responses to light. Light Res Technol 55, 474–486 (2023). 10.1177/14771535221147140

41 Zauner, J., Hartmeyer, S. & Spitschan, M. LightLogR: Reproducible analysis of personal light exposure data. J Open Source Softw 10, 7601 (2025). 10.21105/joss.07601

42 Danilenko, K. V., Stefani, O., Voronin, K. A., Mezhakova, M. S., Petrov, I. M., Borisenkov, M. F., Markov, A. A. & Gubin, D. G. Wearable Light-and-Motion Dataloggers for Sleep/Wake Research: A Review. Appl Sci 12, 11794 (2022). 10.3390/app122211794

43 Guidolin, C., Zauner, J., Hartmeyer, S. L. & Spitschan, M. Collecting, detecting and handling non-wear intervals in longitudinal light exposure data. bioRxiv, 2024.2012.2023.627604 (2024). 10.1101/2024.12.23.627604

44 Bocanegra, G., Stefani, O., Zauner, J., Spitschan, M. & Joosten-Ma, H. Wearable Light Loggers and Dosimeters for Health Impact Measurement: A User Experience Study. SSRN 5047905, (2024). 10.2139/ssrn.5047905

45 Zauner, J., Stefani, O., Bocanegra, G., Guidolin, C., Schrader, B., Udovicic, L. & Spitschan, M. Auxiliary data questionnaires. Zenodo 17535086, (2025). 10.5281/zenodo.17535086

46 Carney, C. E., Buysse, D. J., Ancoli-Israel, S., Edinger, J. D., Krystal, A. D., Lichstein, K. L. & Morin, C. M. The consensus sleep diary: standardizing prospective sleep self-monitoring. Sleep 35, 287–302 (2012). 10.5665/sleep.1642

47 Bajaj, A., Rosner, B., Lockley, S. W. & Schernhammer, E. S. Validation of a light questionnaire with real-life photopic illuminance measurements: the Harvard Light Exposure Assessment questionnaire. Cancer Epidemiol Biomarkers Prev 20, 1341–1349 (2011). 10.1158/1055-9965.EPI-11-0204

48 Zauner, J., Tsukimori, E. & Spitschan, M. LightLogR Newsletter, Edition #01. Zenodo 16308351, (2025). 10.5281/zenodo.16308351

49 Zauner, J., Biller, A. M. & Spitschan, M. Impact of light logger and dosimeter placement on wearability and appeal in real-life settings. Open Res Eur 5, 293 (2025). 10.12688/openreseurope.20985.1

50 Zauner, J., Stefani, O., Biller, A. M., Guidolin, C. & Spitschan, M. Web-based specification tool for wearable light loggers and optical radiation dosimeters (Version 1.0.1) [Software]. Zenodo 1.04ga-fd22, (2025). 10.17617/1.04ga-fd22

51 Murukesu, R. R., Zauner, J. & Spitschan, M. A day in daylight. Zenodo 17467253, (2025). 10.5281/zenodo.17467253

52 Hartmeyer, S. L. & Andersen, M. Towards a framework for light-dosimetry studies: Quantification metrics. Light Res Technol 56, 337–365 (2023). 10.1177/14771535231170500

