## Supplementary material for "Auxiliary data, quality assurance and quality control for wearable light loggers and optical radiation dosimeters": Figure 1 generation

### Supplement 2: Figure1

This file creates Figure 1 of the main paper.

Data available from: https://github.com/tscnlab/ZaunerEtAl_npjBiolTimingSleep_2026/figure/

```r
library(tidyverse)
library(readxl)
library(cowplot)
library(patchwork)
library(mgcv)
library(itsadug)
library(gratia)
```

#### Panel A

```r
#from pubmed
path <- "PubMed_Timeline_Results_by_Year.csv"
data <- read_csv(path, skip = 1)
```

```
Rows: 25 Columns: 2
── Column specification
────────────────────────────────────────────────────────
Delimiter: ","
dbl (2): Year, Count

ℹ Use `spec()` to retrieve the full column specification for this data.
ℹ Specify the column types or set `show_col_types = FALSE` to quiet this
message.
```

```r
#model of publications per year
m1 <- gam(Count ~ s(Year, bs = "tp"), family = poisson, data = data)
m1 |> summary()
```

```


Family: poisson
Link function: log

Formula:
Count ~ s(Year, bs = "tp")

Parametric coefficients:
```

```
            Estimate Std. Error z value Pr(>|z|)
(Intercept)   1.3533     0.1067   12.69   <2e-16 ***
---
Signif. codes:  0 '***' 0.001 '**' 0.01 '*' 0.05 '.' 0.1 ' ' 1

Approximate significance of smooth terms:
          edf Ref.df Chi.sq  p-value
s(Year) 2.144  2.677  29.98 2.52e-06 ***
---
Signif. codes:  0 '***' 0.001 '**' 0.01 '*' 0.05 '.' 0.1 ' ' 1

R-sq.(adj) =  0.546   Deviance explained = 55.3%
UBRE = 0.2212  Scale est. = 1         n = 25
```

```
performance::check_model(m1)
```

```
Cannot simulate residuals for models of class `gam`. Please try
  `check_model(..., residual_type = "normal")` instead.
```

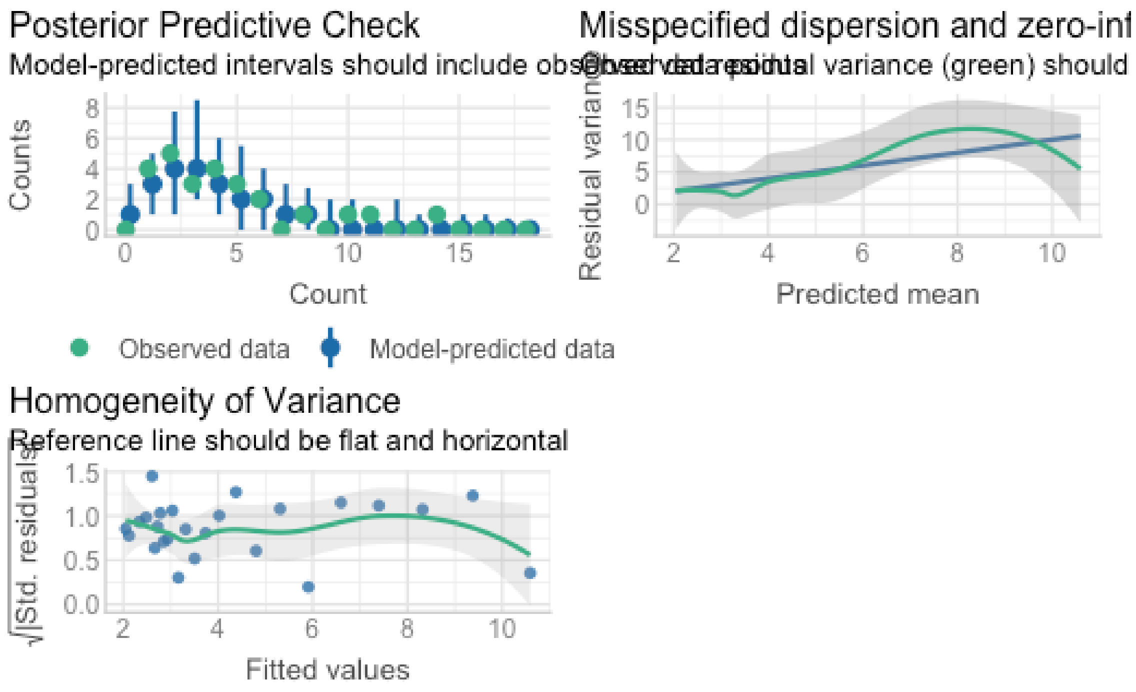


```
draw(m1)
```

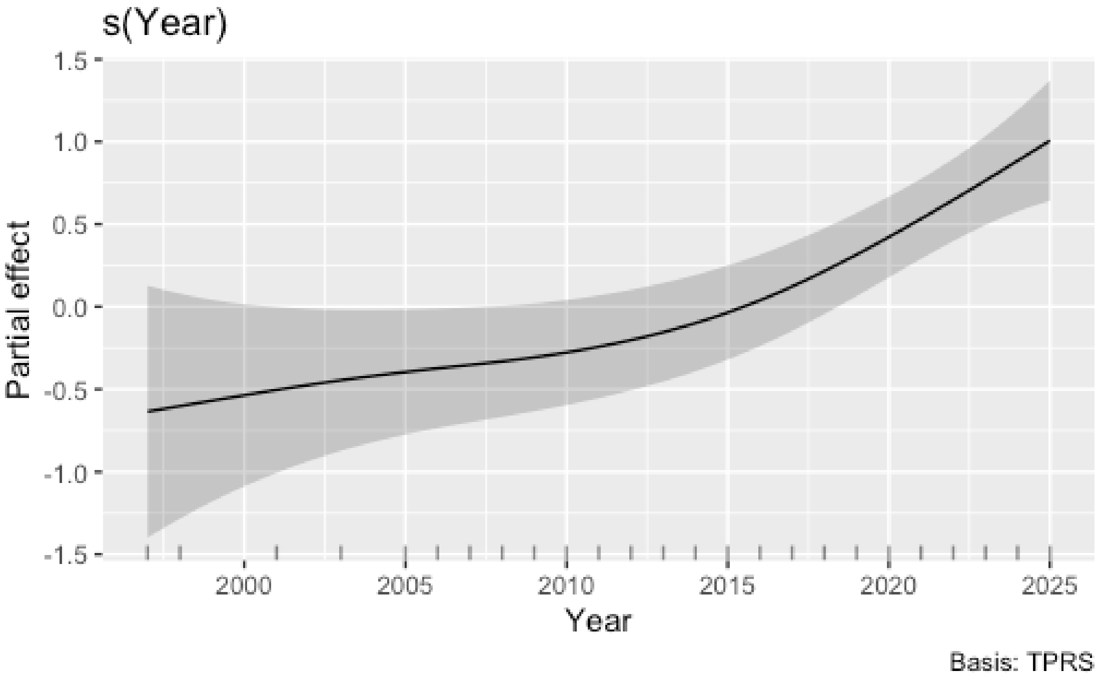


```
#calculating the first derivative of the the smooth
m1d <- derivatives(m1, type = "central")
#calculating the range where the 1st derivative is significantly different
from 0
significant_range <-
m1d |>
  filter(.lower_ci >= 0) |>
  pull(Year) |>
  range() |>
  round()
#figure generation
Fig1a <-
data |>
  ggplot(aes(x=Year, y = Count)) +
  geom_col(fill = "black") +
  theme_cowplot() +
  annotate(geom = "line", x = significant_range, y = -0.65, col = "red", lwd =
1.5) +
  annotate(geom = "point",
           x = significant_range[1],
           y = -0.5, fill = "red", col = "red", size = 4,
           shape = 17)+
  annotate(geom = "text", y = -0.5, x = significant_range[1]+1,
           label = "significant increase",
           col = "red", hjust = 0, vjust = 0, size = 3.25
```

```
          )+
  scale_x_continuous(breaks = c(1997, 2000, 2010, 2014, 2020, 2025)) +
  scale_y_continuous(breaks = c(0, 5, 10, 15)) +
  labs(y = "Light logger publications") +
  coord_cartesian(ylim = c(0, NA), clip = FALSE)
Fig1a
```

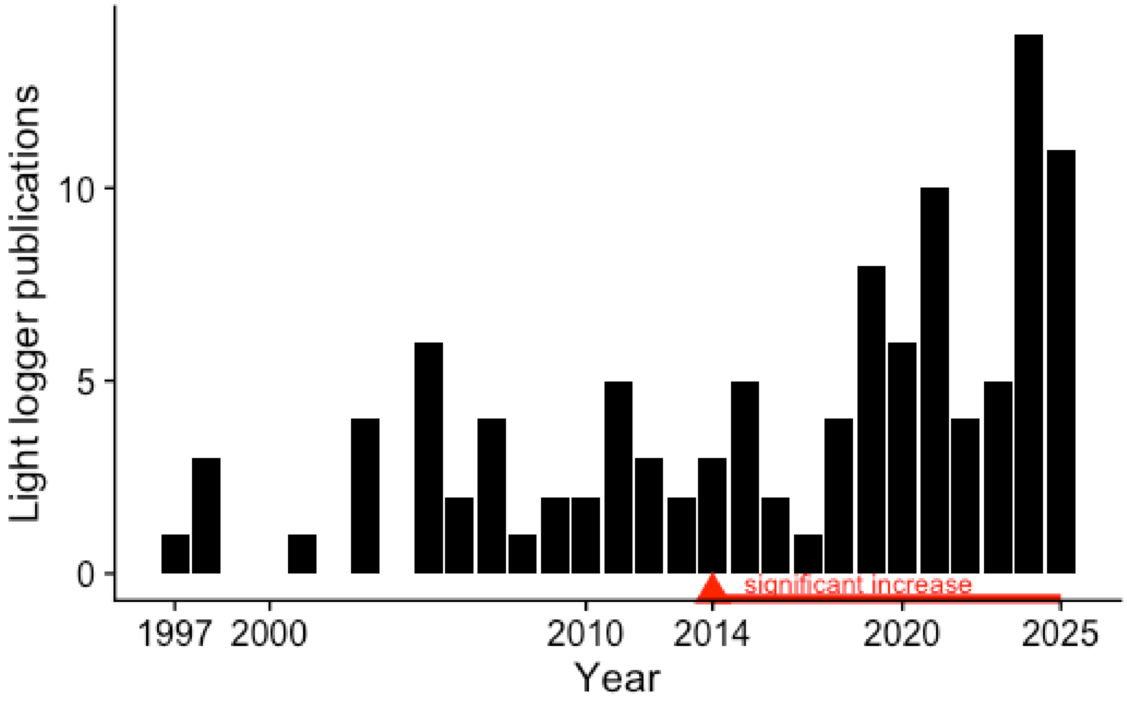


#### Panel B

```
path <- "data_devices.csv"
data_devices <- read_csv(path)
```

```
Rows: 17 Columns: 3
── Column specification
────────────────────────────────────────────────────────
Delimiter: ","
dbl (3): release, n, n_cum

ℹ Use `spec()` to retrieve the full column specification for this data.
ℹ Specify the column types or set `show_col_types = FALSE` to quiet this
message.
```

```
Fig1b <-
data_devices |>
  mutate(n_cum = cumsum(n)) |>
  ggplot(aes(x=release, y = n_cum)) +
  geom_line(lwd = 1.5) +
  scale_x_continuous(breaks = c(2006, 2010, 2015, 2020, 2023)) +
  cowplot::theme_cowplot() +
  labs(x = "Year", y = "Number of light logger models") +
  coord_cartesian(ylim = c(0, 55))

Fig1b
```

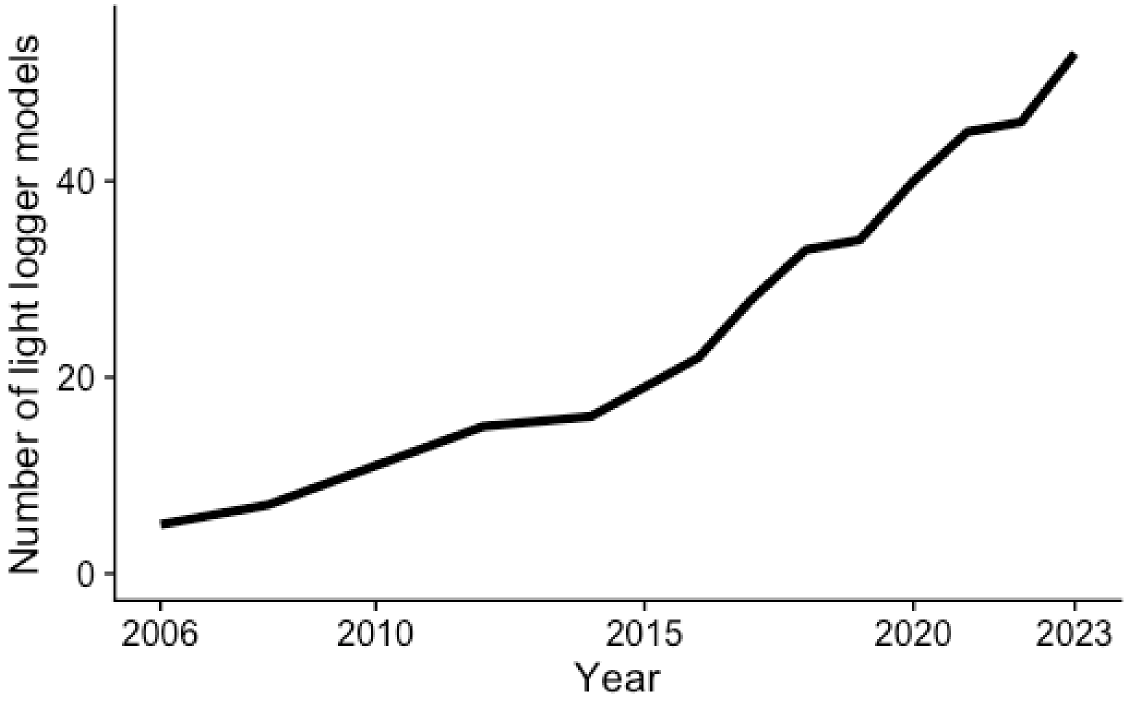


```
Fig1a + Fig1b + plot_annotation(tag_level = "A")
```

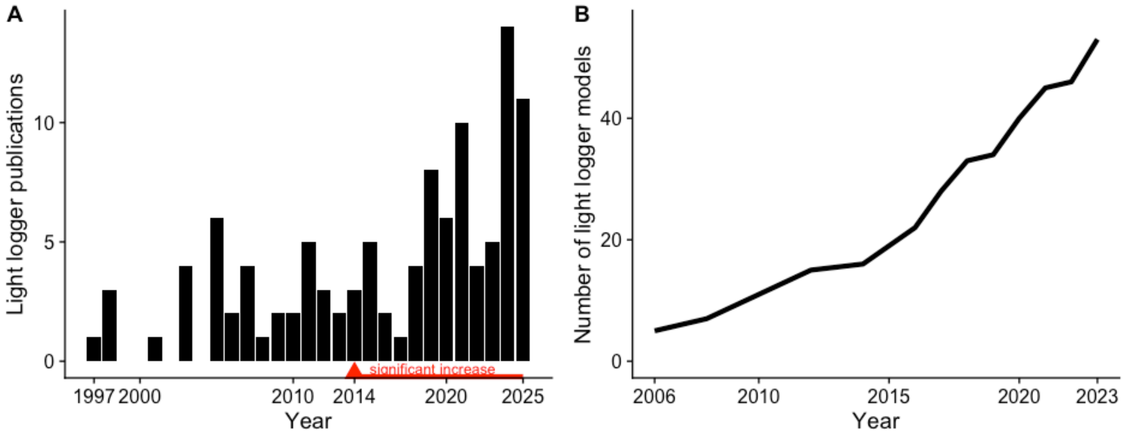


```r
ggsave("Figure_1.png", width = 10, height = 4)

#standalone
Fig1a + Fig1b +
  plot_annotation(tag_level = "A", caption = "Sources: (A) PubMed search for
original research on humans, change analysis with GAM; (B) from van Duijnhoven
et al. (2025)")
```

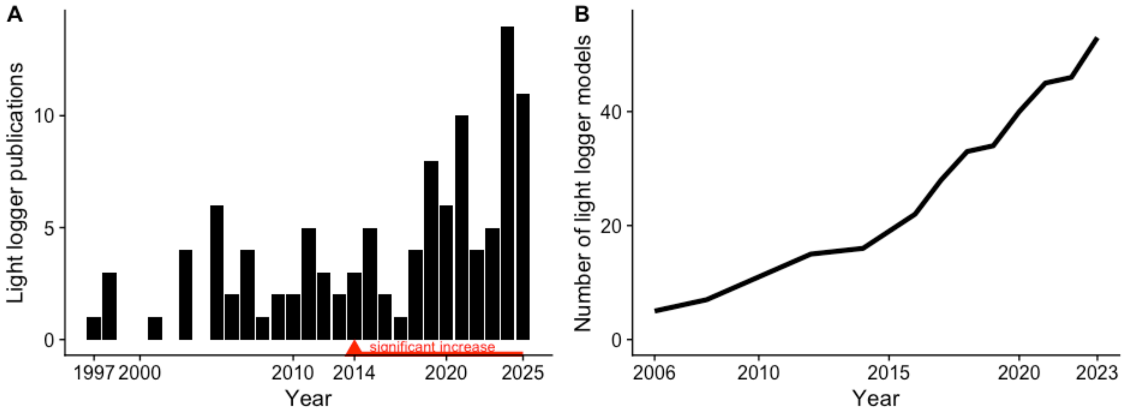


Sources: (A) PubMed search for original research on humans, change analysis with GAM; (B) from van Duijnhoven et al. (2025)

```r
ggsave("Figure_1SA.png", width = 10, height = 4)
```
