## Supplementary material for "Auxiliary data, quality assurance and quality control for wearable light loggers and optical radiation dosimeters": Survey results

### Q1 Do you agree with the definition of auxiliary data as described in the document?

Answered: 15 Skipped: 1

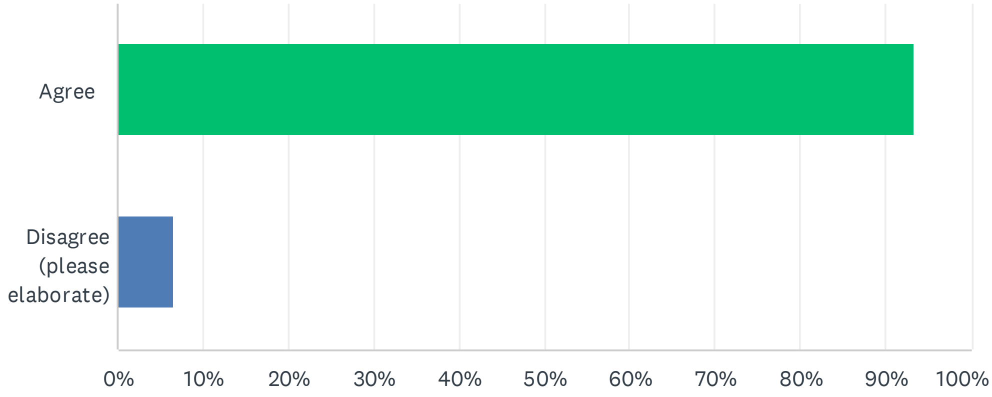


| ANSWER CHOICES | RESPONSES | |
|---|---|---|
| Agree | 93.33% | 14 |
| Disagree (please elaborate) | 6.67% | 1 |
| TOTAL | | 15 |

| # | DISAGREE (PLEASE ELABORATE) | DATE |
|---|---|---|
| 1 | Overall yes, but it is not completely clear to me why the auxiliary data with respect to the user experience is needed (with a high measurement resolution) and how to use this data to invalidate specific assessments. | 5/17/2024 3:44 PM |

### Q2 How have you collected these data previously?

Answered: 16 Skipped: 0

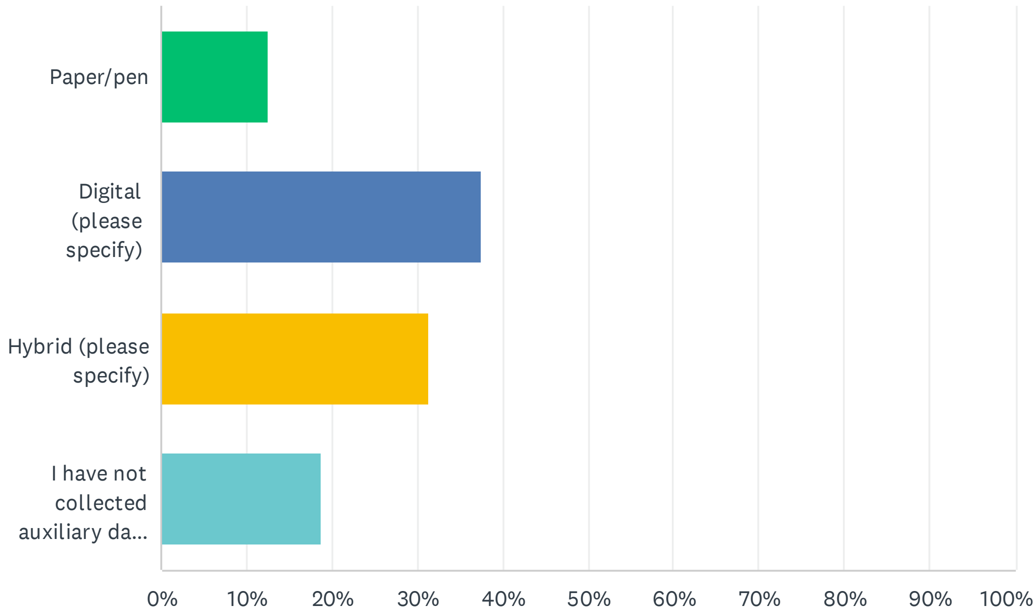


| ANSWER CHOICES | RESPONSES | |
|---|---|---|
| Paper/pen | 12.50% | 2 |
| Digital (please specify) | 37.50% | 6 |
| Hybrid (please specify) | 31.25% | 5 |
| I have not collected auxiliary data before | 18.75% | 3 |
| TOTAL | | 16 |

| # | IF DIGITAL OR HYBRID, PLEASE SPECIFY | DATE |
|---|---|---|
| 1 | I have had patients add to sleep diaries whether or not they used glasses and had light exposure to the agreed upon time. | 6/17/2024 3:04 PM |
| 2 | We generally prefer digital data collection to avoid the added step of manual data entry; however, paper and pencil assessment have sometimes been necessary, in high security environments, when we wanted to avoid additional light from screens, and/or when electronic devices are not available. | 6/13/2024 4:01 PM |
| 3 | Auxiliary data was collected with online questionnaires (redcap. The links for the morning and evening questionnaires were sent to the participants in an automated way with a delay after they filled in the previous one. | 6/11/2024 1:24 PM |
| 4 | We used the REDCap platform to collect the data. However, it involved a general sleep diary questionnaire that asked about bedtime, sleep time, nap time, daylight exposure (time and duration), and activity (time and duration). | 6/11/2024 12:34 PM |
| 5 | I have collected it both in paper forms (alongside a paper sleep diary), and digitally (in Redcap) - but not a mix within the same study, and all studies now use digital collection only. | 6/11/2024 12:18 AM |
| 6 | Using markers on watches to indicate specific periods, including sleep/wake or prolonged off- | 6/10/2024 5:41 PM |

| | | |
|---|---|---|
| | wrist. Similarly, we now sometimes rely on skin temperature data to assess off-wrist | |
| 7 | digital (specific sensors) and online questionnaires filled by participants | 6/6/2024 1:37 PM |
| 8 | We asked the participants to press a trigger button each time they put down the logger and take a note, why they put it down. Additionally, we cleaned the logged data by removing values when there was no movement (movement was recorded by the device). We double checked trigger events, notes and non-movement times before we cleaned the data. | 5/24/2024 1:08 PM |
| 9 | Using an experience sampling application on a smartphone | 5/17/2024 3:44 PM |
| 10 | 1. Environmental light data: I have collected this using a light logger placed on the rooftop of a building. The set-up was designed to mimic that of an optical pyrometer. 2. Activities and light environment throughout the day: I have collected this using pen and paper (from participants). They had to fill this in daily and upload a picture to a shared folder where I could check data quality. Participant entries were then digitalized. 3. Non-wear time: I collected this data using a digital questionnaire that participants could complete on their smartphones. | 5/2/2024 4:03 PM |
| 11 | Study-specific app was designed and used or with Qualtrics data collection online survey services | 5/2/2024 2:30 PM |
| 12 | MyCAP smartphone application | 5/2/2024 2:01 PM |

### Q3 What do you think is the most effective way for collecting these data?

Answered: 16 Skipped: 0

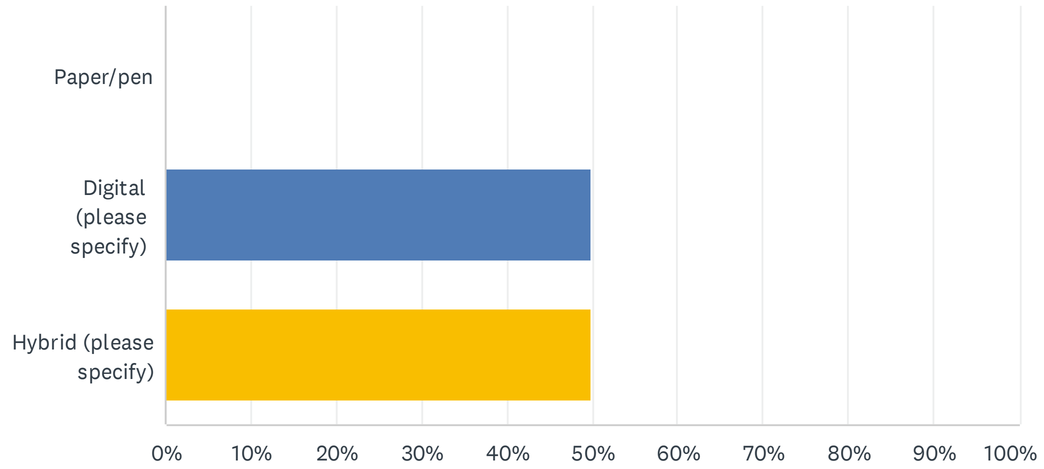


| ANSWER CHOICES | RESPONSES | |
|---|---|---|
| Paper/pen | 0.00% | 0 |
| Digital (please specify) | 50.00% | 8 |
| Hybrid (please specify) | 50.00% | 8 |
| TOTAL | | 16 |

| # | IF DIGITAL OR HYBRID, PLEASE SPECIFY | DATE |
|---|---|---|
| 1 | I think the suggestions in the document was good. | 6/17/2024 3:04 PM |
| 2 | Digital is <generally> easier for everyone, when possible. Qualitative information/open-ended questions may be better collected via paper/pen and/or interview. | 6/13/2024 4:01 PM |
| 3 | Entering data by hand is a large potential source of errors and requires extra workforce resources. Additionally, paper/pencil data can easily get lost. With exceptions, where it is not possible or inconvenient to collect digital data, I prefer collecting these data digitally. | 6/11/2024 1:24 PM |
| 4 | I think using cellphones would be the easiest way to collect data since almost everyone uses them. However, it's always best to give participants the option to choose what works best for them, as some people might prefer a paper version. | 6/11/2024 12:34 PM |
| 5 | I use RedCap for ease of data organisation, specifically my preference is the MyCap app as I find this promotes the best compliance. | 6/11/2024 12:18 AM |
| 6 | I think a combination of marker usage + paper/pen would be a complete way of assessing. Sometimes, participants do not have a way of noting things down and do want to mark - so digital marking is a must | 6/10/2024 5:41 PM |
| 7 | Participants tend to underestimate the auxiliary data so its needed to have wearing sensors on teh device but sometimes classification can be hard and so adding a questionnaire is useful. | 6/6/2024 1:37 PM |
| 8 | Digigital data such as movement, temperature, skin conductance will make the process most effective but a double check with manual notes from the participant will ensure confidence with the measured light levels. Example: the device could deliver movement data when it was put into a bag while moving. | 5/24/2024 1:08 PM |

| | | |
|---|---|---|
| 9 | I don't think there is a way of collecting these data that will be effective across all study populations. Researchers should evaluate relevant characteristics of individuals such as age, cognitive limitations, technological proficiency, and daily routines, as these can affect their adherence to data collection tools. Thus, tailoring the data collection strategy to the specific needs and capabilities of the study population is important. This can involve gathering feedback from participants and being flexible in adapting the methods used. | 5/17/2024 4:15 PM |
| 10 | Allows you to have time-stamped data and do real-time monitoring of data collection. The choice for the modality might also depend on the target population. | 5/17/2024 3:44 PM |
| 11 | Some data are easier to be record automatically with digital. While some volunteers may prefer paper works. | 5/12/2024 5:17 PM |
| 12 | Either through SMS or SMS-based links. Can confirm compliance much more easily. | 5/2/2024 11:54 PM |
| 13 | I think it depends on the individual auxiliary data. In the categories that you have listed: - Wear/non-wear: digital questionnaire would be best - Sleep/wake: digital questionnaire - Light environment throughout the day: it depends on how this data is collected. If every time participants change light environment, they update this in real time, then I would suggest that this should be in the form of a "digital log". However, if this is a retrospective questionnaire (e.g. participants filling in, at the end of their day, what type of light exposure they had for every hour of the day), then a paper form would be best to visualise this. The reason is that you can visualise time chronologically (so 2PM comes after 1PM), which helps individuals reconstruct their days and locations, and thus light environments, better than asking the question "What light were you exposed to at 2PM?". - Behaviour/exercise: same as above - Experience with the light logger: digital questionnaire - Environmental light levels: possibly with a calibrated device which measures light continuously as participants take part in the experiment. If this is not possible, a representative measurement of light at specific times, such as sunset, during data collection would be useful. If this is also not available, photoperiod information could be retrieved by climate tools available online. | 5/2/2024 4:03 PM |
| 14 | Study-specific app with reminders to data entry | 5/2/2024 2:30 PM |
| 15 | Any ESM application such as MyCAP, in which questionnaires can be timed and sent to the participants. Ability to send reminders to fill in pending questionnaires, as well as retrospective questionnaire completion is very helpful. | 5/2/2024 2:01 PM |

### Q4 How important do you think auxiliary data as defined is in general for any light exposure data collection effort?

Answered: 16 Skipped: 0

4.0
average rating

| | ISN'T REQUIRED OR HELPFUL FOR MY TYPE OF RESEARCH | (NO LABEL) | (NO LABEL) | (NO LABEL) | LIGHT EXPOSURE MEASUREMENT WOULD BE WORTHLESS WITHOUT IT | TOTAL | WEIGHTED AVERAGE |
|---|---|---|---|---|---|---|---|
| ☆ | 0.00%<br>0 | 0.00%<br>0 | 18.75%<br>3 | 62.50%<br>10 | 18.75%<br>3 | 16 | 4.00 |

### Q5 Which domain of auxiliary data are you most interested in / do you think are relevant to your research? (Order them by relevance or discard them with the checkbox)

Answered: 15 Skipped: 1

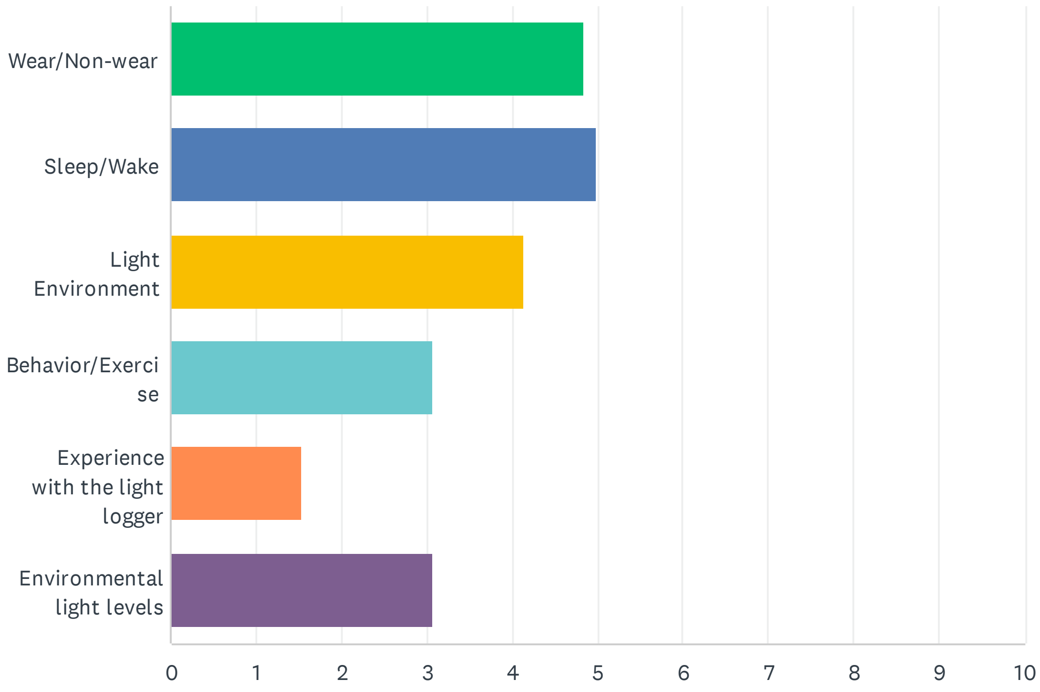


| | 1 | 2 | 3 | 4 | 5 | 6 | NOT RELEVANT | TOTAL | SCORE |
|---|---|---|---|---|---|---|---|---|---|
| Wear/Non-wear | 26.67%<br>4 | 20.00%<br>3 | 40.00%<br>6 | 0.00%<br>0 | 0.00%<br>0 | 0.00%<br>0 | 13.33%<br>2 | 15 | 4.85 |
| Sleep/Wake | 46.67%<br>7 | 26.67%<br>4 | 0.00%<br>0 | 13.33%<br>2 | 6.67%<br>1 | 0.00%<br>0 | 6.67%<br>1 | 15 | 5.00 |
| Light Environment | 13.33%<br>2 | 20.00%<br>3 | 40.00%<br>6 | 20.00%<br>3 | 6.67%<br>1 | 0.00%<br>0 | 0.00%<br>0 | 15 | 4.13 |
| Behavior/Exercise | 6.67%<br>1 | 13.33%<br>2 | 13.33%<br>2 | 26.67%<br>4 | 26.67%<br>4 | 13.33%<br>2 | 0.00%<br>0 | 15 | 3.07 |
| Experience with the light logger | 0.00%<br>0 | 0.00%<br>0 | 0.00%<br>0 | 6.67%<br>1 | 33.33%<br>5 | 46.67%<br>7 | 13.33%<br>2 | 15 | 1.54 |
| Environmental light levels | 6.67%<br>1 | 20.00%<br>3 | 6.67%<br>1 | 26.67%<br>4 | 20.00%<br>3 | 20.00%<br>3 | 0.00%<br>0 | 15 | 3.07 |

### Q6 Are there other domains of auxiliary data as defined that you think are important yet missing? (please elaborate)

Answered: 10 Skipped: 6

| # | RESPONSES | DATE |
|---|---|---|
| 1 | where on the body the light sensor is being worn (for devices where that is an option and/or may vary for participant comfort); information about accessories that would alter light entering the eyes (e.g. hats, visors, blue blockers, sleep masks, etc) | 6/13/2024 4:01 PM |
| 2 | Depending on where the light logger is worn, data on clothing could be interesting. | 6/11/2024 1:24 PM |
| 3 | However, we always instruct participants not to cover the sensor with sleeves. Still, we could include questions about long sleeves, such as their color and texture. | 6/11/2024 12:34 PM |
| 4 | Perhaps more relevant for loggers which are not glasses based, but wear data for typical glasses/sunglasses may be relevant, and potential coverage (though this can usually be ascertained from the data its self). | 6/11/2024 12:18 AM |
| 5 | Reasons for not wearing a device | 5/24/2024 1:08 PM |
| 6 | I think the draft version of the auxiliary data strategy provided is very complete and well-thought. I would suggest minor points: 1) If relevant to the research project, I would suggest asking participants to better describe their indoor light environment. For example asking them about curtains (do they sleep with closed curtains? do they keep curtains opened during the day? And so on). 2) In the "experience log" section, I missed a question specifying what prevented the participant from using the light glasses/light wearables (aesthetic issues? non-adherence due to forgetfulness? other reasons?). This could help researchers refine data collection for future studies. 3) I think there should be a new section of auxiliary data for light intervention studies (especially if conducted in real-world settings). This section could include questions about participants' expectations and acceptability of the intervention, any symptoms or side-effects caused by the light intervention, times and durations of light intervention exposure, overall participant experience/feedback on the intervention. | 5/17/2024 4:15 PM |
| 7 | no | 5/12/2024 5:17 PM |
| 8 | This would be dependent on the form factor of the device. This seems written for a glasses-based device and it is unclear whether the accuracy of this outweighs the fashion/inconvenience aspect in field studies. | 5/2/2024 11:54 PM |
| 9 | 1) Distinguishing work day and free day (not just weekday and weekends. Some sleep diaries do this but not all). 2) Understanding the profession of the individual wearing the light logger. 3) Reporting temperature. In some seasons or certain countries, ambient temperature might determine whether someone stays indoors or outdoors (if it is 37 degrees, there is a chance I will have my lunch break indoors rather than outdoors because too warm). 4) Perhaps also report weather conditions (an addition to the environmental light logging) | 5/2/2024 4:03 PM |
| 10 | This structure may not work for irragular sleepers or night shift workers, as some surveys e.g. sleep diary is designed to capture sleep once in a day after mornign wake | 5/2/2024 2:30 PM |

### Q7 How important do you think this domain is?

Answered: 16 Skipped: 0

3.6★

average rating

|  | ISN'T REQUIRED OR HELPFUL FOR MY TYPE OF RESEARCH | (NO LABEL) | (NO LABEL) | (NO LABEL) | LIGHT EXPOSURE MEASUREMENT WOULD BE WORTHLESS WITHOUT IT | TOTAL | WEIGHTED AVERAGE |
|---|---|---|---|---|---|---|---|
| ☆ | 6.25%<br>1 | 6.25%<br>1 | 25.00%<br>4 | 43.75%<br>7 | 18.75%<br>3 | 16 | 3.63 |

### Q8 How well do you think the suggested measures and procedures are able to capture the domain?

Answered: 16 Skipped: 0

4.1 ★

average rating

| | NOT AT ALL | NOT WELL | PARTLY | SUFFICIENTLY | COMPLETELY | TOTAL | WEIGHTED AVERAGE |
|---|---|---|---|---|---|---|---|
| ☆ | 0.00%<br>0 | 0.00%<br>0 | 6.25%<br>1 | 75.00%<br>12 | 18.75%<br>3 | 16 | 4.13 |

| # | IF 3 STARS OR LESS: DO YOU HAVE SUGGESTIONS FOR IMPROVEMENT? | DATE |
|---|---|---|
| 1 | I'm not sure how to answer this question. I think the log would provide useful information to have but I am skeptical about participants filling it all out in a meaningful way, especially if they have other study activities they need to complete. I think it is perhaps too burdensome/detailed. The event button is nice, though not all photosensors contain one. I also think a single field could be added to the sleep diary for participants to indicated approximate times of removal. | 6/13/2024 4:06 PM |
| 2 | I think the participant instructions and burden are very critical here - at the moment there are quite a number of questionnaires, and if participants were asked to log anything longer than 1 minute, answering several questions each time, I think compliance could be concern (but of course this is dependent on how many things other than light logging the participant is being asked to complete!). If it was this set of questionnaires alone, I think it would be feasible/effective - but for most studies I would be doing, I would try and simplify the questions here (e.g., just recording the time the logger was put back on, and then estimate time off, as usually it is not too tricky to infer a non-wear period in the data if you have an approximate time period). Largely though, as this is so dependent on compliance, I think the questionnaire does as good a job as possible if we assume people complete it fully each time. | 6/11/2024 12:28 AM |
| 3 | I would add a 6th option to the Wear Log questionnaire: - "Putting the light glasses on upon waking up" | 5/2/2024 4:05 PM |

### Q9 If you want to give additional feedback on this domain, please do so here

Answered: 5 Skipped: 11

| # | RESPONSES | DATE |
|---|---|---|
| 1 | The way this questionnaire is set up makes it difficult to address practicality, which is an important piece of the puzzle. | 6/13/2024 4:06 PM |
| 2 | The proposed assessment seems to be rather cumbersome for the participants. Sensor data to detect wearing/not wearing might be preferred | 5/17/2024 3:45 PM |
| 3 | Not yet. | 5/12/2024 5:20 PM |
| 4 | Would seem this is better captured on device as remembering to fill this out each time could be a considerable burden. Why not just wear/non-wear? Could be logged easier. | 5/2/2024 11:56 PM |
| 5 | I believe that well validated automated algorithms for the identification of non-wear periods trump any non-wear log, since it cannot be excluded that participants forget to perform the suggested measures and then algorithms like this have to be employed anyways. Moreover, well constructed algorithms should be able to identify most of the invalid data due to non-wear. The advantage of the non-wear log as proposed here is the documentation of the location and activity of participants during the non-wear period. | 5/2/2024 2:07 PM |

### Q10 How important do you think this domain is?

Answered: 16 Skipped: 0

4.0

average rating

| | ISN'T REQUIRED OR HELPFUL FOR MY TYPE OF RESEARCH | (NO LABEL) | (NO LABEL) | (NO LABEL) | LIGHT EXPOSURE MEASUREMENT WOULD BE WORTHLESS WITHOUT IT | TOTAL | WEIGHTED AVERAGE |
|---|---|---|---|---|---|---|---|
| ☆ | 0.00%<br>0 | 12.50%<br>2 | 12.50%<br>2 | 37.50%<br>6 | 37.50%<br>6 | 16 | 4.00 |

### Q11 How well do you think the suggested measures and procedures are able to capture the domain?

Answered: 16 Skipped: 0

4.0

average rating

| | NOT AT ALL | NOT WELL | PARTLY | SUFFICIENTLY | COMPLETELY | TOTAL | WEIGHTED AVERAGE |
|---|---|---|---|---|---|---|---|
| ☆ | 0.00%<br>0 | 6.25%<br>1 | 18.75%<br>3 | 43.75%<br>7 | 31.25%<br>5 | 16 | 4.00 |

| # | IF 3 STARS OR LESS: DO YOU HAVE SUGGESTIONS FOR IMPROVEMENT? | DATE |
|---|---|---|
| 1 | A lot of times, participants will be using photosensors with actigraphy, and those more objective sleep parameters are likely to be preferrable. One issue with this is that if two separate devices are needed, clocks for the devices must be synchronized. Still, it is more precise in our experience than diary data. But the necessity for precision will depend on the study questions, sample sizes, etc. | 6/13/2024 4:09 PM |
| 2 | Seems redundant with the wear/non-wear. Not clear on what additional data about contextualizing light this is providing. | 5/2/2024 11:57 PM |
| 3 | I would use a Sleep diary which also enables collection of nap times, not just the main sleep episode. | 5/2/2024 4:09 PM |

### Q12 If you want to give additional feedback on this domain, please do so here

Answered: 2 Skipped: 14

| # | RESPONSES | DATE |
|---|---|---|
| 1 | The necessity of this may differ if a validated sleep tracker is being used alongside light logging. | 6/11/2024 12:28 AM |
| 2 | I think it all depends on what the goal of the measurement is. Sleep diaries can be important if interested in timing of sleep/wake, but I tend to use actigraphy independent of this (as it is so inaccurate) | 6/10/2024 5:43 PM |

### Q13 How important do you think this domain is?

Answered: 16 Skipped: 0

3.4

average rating

| | ISN'T REQUIRED OR HELPFUL FOR MY TYPE OF RESEARCH | (NO LABEL) | (NO LABEL) | (NO LABEL) | LIGHT EXPOSURE MEASUREMENT WOULD BE WORTHLESS WITHOUT IT | TOTAL | WEIGHTED AVERAGE |
|---|---|---|---|---|---|---|---|
| ☆ | 6.25%<br>1 | 6.25%<br>1 | 43.75%<br>7 | 31.25%<br>5 | 12.50%<br>2 | 16 | 3.38 |

### Q14 How well do you think the suggested measures and procedures are able to capture the domain?

Answered: 15 Skipped: 1

3.9

average rating

| | NOT AT ALL | NOT WELL | PARTLY | SUFFICIENTLY | COMPLETELY | TOTAL | WEIGHTED AVERAGE |
|---|---|---|---|---|---|---|---|
| ☆ | 0.00%<br>0 | 6.67%<br>1 | 26.67%<br>4 | 40.00%<br>6 | 26.67%<br>4 | 15 | 3.87 |

| # | IF 3 STARS OR LESS: DO YOU HAVE SUGGESTIONS FOR IMPROVEMENT? | DATE |
|---|---|---|
| 1 | This is another case where I think it is a lot of information to expect participants to provide. I don't know that the data will be meaningful and if the burden of collection is worth the information gleaned. If all you are collecting is light data, then I think all of this is fine. But if light data is just one of several measures, I think this level of input is too much. I would also make the suggestion, for all of these strategies, that a reminder system be in place, which can increase compliance and quality of data. Another piece of information that needs to be clarified here is how should all of this data be integrated. Like, what it the data from the light exposure dairy conflicts with the wear log. What is the hierarchy you would employ? | 6/13/2024 4:14 PM |
| 2 | For now, the idea of specifying different light exposures is great, but my concern is that participants often complain about filling out too many log books. In the future, perhaps we could include buttons on the logger itself, allowing participants to simply press a button to record their light exposure. This would likely make the process easier and more user-friendly. | 6/11/2024 12:35 PM |
| 3 | This captures exposure very thoroughly, but I think for many use cases it would be beyond what is feasible unless light is the primary/sole outcome. | 6/11/2024 12:28 AM |
| 4 | The relative contribution of these in many environments would be unknowable by the participant (in a classroom with a window, overhead lighting, and a laptop - how much of each source is impacting the light reaching the cornea? Seems like more information that is useful in most circumstances. | 5/2/2024 11:58 PM |
| 5 | Suggest splitting this into two questionnaires, each covering 12 hours, and having participants fill this in twice a day (i.e. at lunch time and before sleep). This would make it easier for them to recall their light environments. | 5/2/2024 4:12 PM |

### Q15 If you want to give additional feedback on this domain, please do so here

Answered: 4 Skipped: 12

| # | RESPONSES | DATE |
|---|---|---|
| 1 | Also here, I think it depends on the utility. If using time above threshold, the actiwatches are quite accurate and then the light diary becomes less important. | 6/10/2024 5:44 PM |
| 2 | The diary helps for the data analysis | 6/6/2024 1:44 PM |
| 3 | I think that the relevance is largely dependent on the research question | 5/17/2024 3:46 PM |
| 4 | I think the quality of this data and gains made from it do not outweigh the participant burden of having to fill in this additional log. | 5/2/2024 2:08 PM |

### Q16 How important do you think this domain is?

Answered: 16 Skipped: 0

3.2 ★

average rating

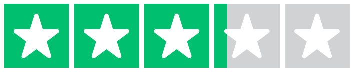

| | ISN'T REQUIRED OR HELPFUL FOR MY TYPE OF RESEARCH | (NO LABEL) | (NO LABEL) | (NO LABEL) | LIGHT EXPOSURE MEASUREMENT WOULD BE WORTHLESS WITHOUT IT | TOTAL | WEIGHTED AVERAGE |
|---|---|---|---|---|---|---|---|
| ☆ | 6.25%<br>1 | 12.50%<br>2 | 43.75%<br>7 | 31.25%<br>5 | 6.25%<br>1 | 16 | 3.19 |

### Q17 How well do you think the suggested measures and procedures are able to capture the domain?

Answered: 16 Skipped: 0

4.1 ★
average rating

| | NOT AT ALL | NOT WELL | PARTLY | SUFFICIENTLY | COMPLETELY | TOTAL | WEIGHTED AVERAGE |
|---|---|---|---|---|---|---|---|
| ☆ | 0.00%<br>0 | 6.25%<br>1 | 12.50%<br>2 | 50.00%<br>8 | 31.25%<br>5 | 16 | 4.06 |

| # | IF 3 STARS OR LESS: DO YOU HAVE SUGGESTIONS FOR IMPROVEMENT? | DATE |
|---|---|---|
| 1 | I feel like this information is a one-off from the light exposure information and may further confuse what is really going on, if it doesn't align with other data being collected. On the plus side, it might also be interesting information for understanding photic exposure during specific activities and where are the most important points of intervention. Again, if you are only collecting light data, it is probably worth collection; however, if you have several other assessments, this feels like more burden on participants than it is worth. I do like the confidence scales that accompany the likert scores, which you have elsewhere too, as perhaps this can be used for prioritizing data when it is inconsistent. But it would be helpful to know how you would recommend doing that (e.g. is the data weighted, does the data with most confidence win, etc) | 6/13/2024 4:19 PM |
| 2 | The same as the previous question | 6/11/2024 12:35 PM |
| 3 | A bit of a misnomer (more behavior than exercise). As with the previous lighting question, I am dubious about the probative value of these questions for most research. | 5/3/2024 12:00 AM |

### Q18 If you want to give additional feedback on this domain, please do so here

Answered: 3 Skipped: 13

| # | RESPONSES | DATE |
|---|---|---|
| 1 | Small note - I interpreted exercise as being physical activity/working out, which participants could do also. | 6/11/2024 12:28 AM |
| 2 | The relevance will also be dependent on the research question | 5/17/2024 3:46 PM |
| 3 | I would add sport as a category of its own, and specify if this is performed indoors or outdoors | 5/2/2024 4:13 PM |

### Q19 How important do you think this domain is?

Answered: 16    Skipped: 0

**3.0** ★

average rating

| | ISN'T REQUIRED OR HELPFUL FOR MY TYPE OF RESEARCH | (NO LABEL) | (NO LABEL) | (NO LABEL) | LIGHT EXPOSURE MEASUREMENT WOULD BE WORTHLESS WITHOUT IT | TOTAL | WEIGHTED AVERAGE |
|---|---|---|---|---|---|---|---|
| ☆ | 6.25%<br>1 | 37.50%<br>6 | 25.00%<br>4 | 12.50%<br>2 | 18.75%<br>3 | 16 | 3.00 |

### Q20 How well do you think the suggested measures and procedures are able to capture the domain?

Answered: 16 Skipped: 0

3.8 ★

average rating

| | NOT AT ALL | NOT WELL | PARTLY | SUFFICIENTLY | COMPLETELY | TOTAL | WEIGHTED AVERAGE |
|---|---|---|---|---|---|---|---|
| ☆ | 0.00%<br>0 | 12.50%<br>2 | 25.00%<br>4 | 37.50%<br>6 | 25.00%<br>4 | 16 | 3.75 |

| # | IF 3 STARS OR LESS: DO YOU HAVE SUGGESTIONS FOR IMPROVEMENT? | DATE |
|---|---|---|
| 1 | I think it would be helpful to have less albeit more general items accompanied by one field for more open-ended comments. For example, items about: satisfaction, reason(s) for satisfaction score, ease of use, reasons for ease of use score, ways to improve the experience in the future, etc (all with a list of multiple choice options, including "other" for which they may check all that apply) followed by, "Is there anything else you would like to share with us about your experience." In our experience, multiple choice options for these sorts of things give us more/better data. And if you really wanted to dig into this more, interviews with participants may provide information beyond this. | 6/13/2024 4:29 PM |
| 2 | I think the light logger should be aesthetically pleasing to be worn all day without constraint, making the participant's experience important to improve wearing time. | 6/6/2024 1:47 PM |
| 3 | Standards such as the SUS or WEAR questionnaires are only partly suitable for usability testing of light loggers. | 5/24/2024 1:13 PM |
| 4 | The reporting seems to be a bit complex for the participants | 5/17/2024 3:48 PM |

### Q21 If you want to give additional feedback on this domain, please do so here

Answered: 3 Skipped: 13

| # | RESPONSES | DATE |
| --- | --- | --- |
| 1 | This captures the domain well, but obviously, the utility will depend a little on the research question here. I don't think this is something that is necessary to collect alongside larger research questions (e.g., about associations between light exposure and mood etc), but it is helpful during the validation of a logger or when looking at feasibility for use at scale. | 6/11/2024 12:29 AM |
| 2 | Seems only helpful in limited validation studies. The open-ended nature of the question may lead to inconsistent responses that would be difficult to condense across an array of participants. | 5/3/2024 12:01 AM |
| 3 | Ensure that this is performed on a regular basis, e.g. at least three times per week or similar. Since participants might not feel obliged to complete this, as the instrctions do not define a specific frequency of when they should do so, they might forget about the questionnaire | 5/2/2024 4:15 PM |

### Q22 How important do you think this domain is?

Answered: 15 Skipped: 1

3.7 ★

average rating

| | ISN'T REQUIRED OR HELPFUL FOR MY TYPE OF RESEARCH | (NO LABEL) | (NO LABEL) | (NO LABEL) | LIGHT EXPOSURE MEASUREMENT WOULD BE WORTHLESS WITHOUT IT | TOTAL | WEIGHTED AVERAGE |
|---|---|---|---|---|---|---|---|
| ☆ | 0.00%<br>0 | 6.67%<br>1 | 40.00%<br>6 | 33.33%<br>5 | 20.00%<br>3 | 15 | 3.67 |

### Q23 How well do you think the suggested measures and procedures are able to capture the domain?

Answered: 15 Skipped: 1

3.9★

average rating

| | NOT AT ALL | NOT WELL | PARTLY | SUFFICIENTLY | COMPLETELY | TOTAL | WEIGHTED AVERAGE |
|---|---|---|---|---|---|---|---|
| ☆ | 0.00%<br>0 | 6.67%<br>1 | 20.00%<br>3 | 53.33%<br>8 | 20.00%<br>3 | 15 | 3.87 |

| # | IF 3 STARS OR LESS: DO YOU HAVE SUGGESTIONS FOR IMPROVEMENT? | DATE |
|---|---|---|
| 1 | I think this is a nice idea and easy enough to do. I think the necessity of this really depends on your study and in most cases, it will not be needed. | 6/13/2024 4:31 PM |
| 2 | global illuminance levels from metrological institutions do not precisely re-cap the environmental light levels near participants. Sensors at the roof only provide valuable measurements if the participant did not move into other buildings, etc. | 5/24/2024 1:17 PM |
| 3 | Is a rooftop reflective of anything more (in terms of gross magnitude, not accuracy) than you could get from satellite data (especially given the difficulty of obtaining the data)? Given all of the behavioral and environmental aspects of light (trees, buildings, etc.), is this helpful? | 5/3/2024 12:03 AM |
| 4 | Information about how the sensor should be installed on the rooftop should be indicated. That is, vertical orientation, and if yes, in which cardinal direction? Horizontal orientation? | 5/2/2024 2:13 PM |

### Q24 If you want to give additional feedback on this domain, please do so here

Answered: 2 Skipped: 14

| # | RESPONSES | DATE |
|---|---|---|
| 1 | My only concern here is feasibility, but the described protocol would be a great way to do it! | 6/11/2024 12:34 AM |
| 2 | Position in the room might also be relevant | 5/17/2024 3:48 PM |

### Q25 Do you think the auxiliary data strategy is helpful for the design of your future research?

Answered: 15 Skipped: 1

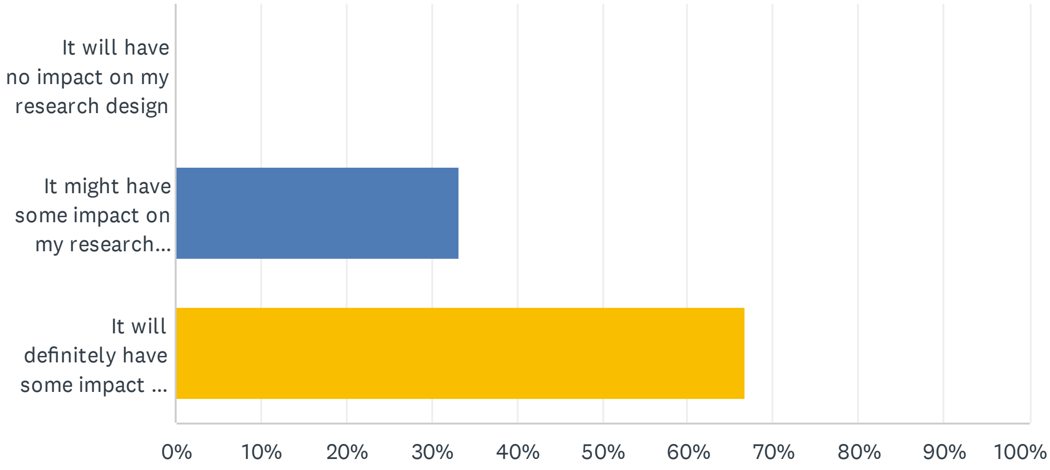


| ANSWER CHOICES | RESPONSES | |
|---|---|---|
| It will have no impact on my research design | 0.00% | 0 |
| It might have some impact on my research design | 33.33% | 5 |
| It will definitely have some impact on my research design | 66.67% | 10 |
| TOTAL | | 15 |

### Q26 If you want to give general feedback on the auxiliary data strategy, please do so here

Answered: 5 Skipped: 11

| # | RESPONSES | DATE |
|---|---|---|
| 1 | I think these sorts of prescriptive things are hard to do because they really depend so much on your study, the population, etc. I've already provided additional general feedback earlier in open fields. | 6/13/2024 4:33 PM |
| 2 | In general, I like the idea of this project. It would definitely help us obtain more reliable data from wearable loggers, making it easier to analyze and interpret the data. Thanks for your effort! | 6/11/2024 12:41 PM |
| 3 | If the intention is to create a strategy which can be applied across studies/research groups, it would be helpful to use either more neutral language, or have options for researchers to select the specific type of logger they are using (glasses, wrist worn, body worn) to have customised question formats. | 6/11/2024 12:38 AM |
| 4 | Sorry to be the person who says "it depends", but I think that I tried to find ways around needing to use auxiliary data (i.e., not relying on sleep/wake timing, only using time above threshold). I do think this can add a lot to the studies and study design | 6/10/2024 5:46 PM |
| 5 | I think auxiliary data are always needed | 6/6/2024 1:48 PM |

### Q27 Would you like to be named as a contributor to the auxiliary data questionnaire in reports and publications about the topic?

Answered: 15 Skipped: 1

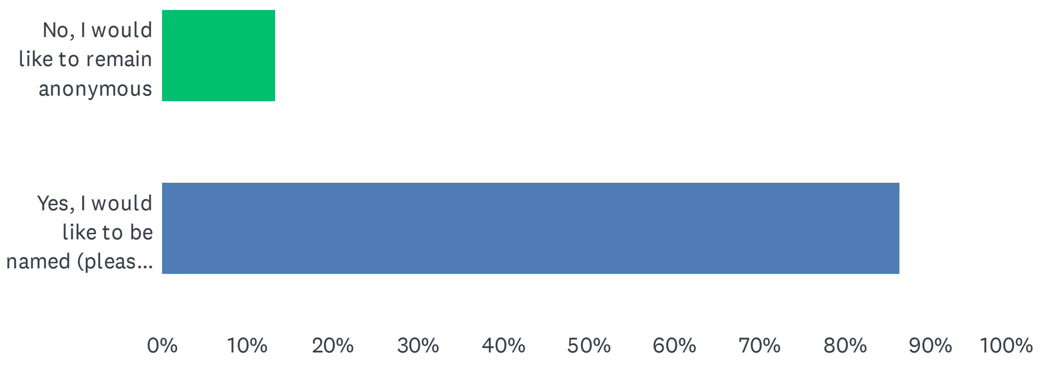


| ANSWER CHOICES | RESPONSES | |
|---|---|---|
| No, I would like to remain anonymous | 13.33% | 2 |
| Yes, I would like to be named (please provide name and email) | 86.67% | 13 |
| TOTAL | | 15 |

| # | YES, I WOULD LIKE TO BE NAMED (PLEASE PROVIDE NAME AND EMAIL) | DATE |
|---|---|---|
| 1 | Gena Glickman [redacted] | 6/13/2024 4:33 PM |
| 2 | Rafael Lazar [redacted] | 6/11/2024 1:30 PM |
| 3 | Fatemeh Fazlali - [redacted] | 6/11/2024 12:41 PM |
| 4 | Elise McGlashan, [redacted] | 6/11/2024 12:38 AM |
| 5 | Renske Lok, [redacted] | 6/10/2024 5:46 PM |
| 6 | Oliver Stefani [redacted] | 5/24/2024 1:18 PM |
| 7 | Débora B. Constantino [redacted] | 5/17/2024 4:17 PM |
| 8 | Karin Smolders; [redacted] | 5/17/2024 3:49 PM |
| 9 | tianren.chen[redacted] | 5/13/2024 1:34 PM |
| 10 | Jamie Zeitzer, [redacted] | 5/3/2024 12:04 AM |
| 11 | Carolina Guidolin, [redacted] | 5/2/2024 4:17 PM |
| 12 | Altug Didikoglu - [redacted] | 5/2/2024 2:34 PM |
| 13 | Steffen Hartmeyer, [redacted] | 5/2/2024 2:13 PM |

### Q28 Would you be willing to be contacted again regarding this topic or the MeLiDos project in general?

Answered: 15 Skipped: 1

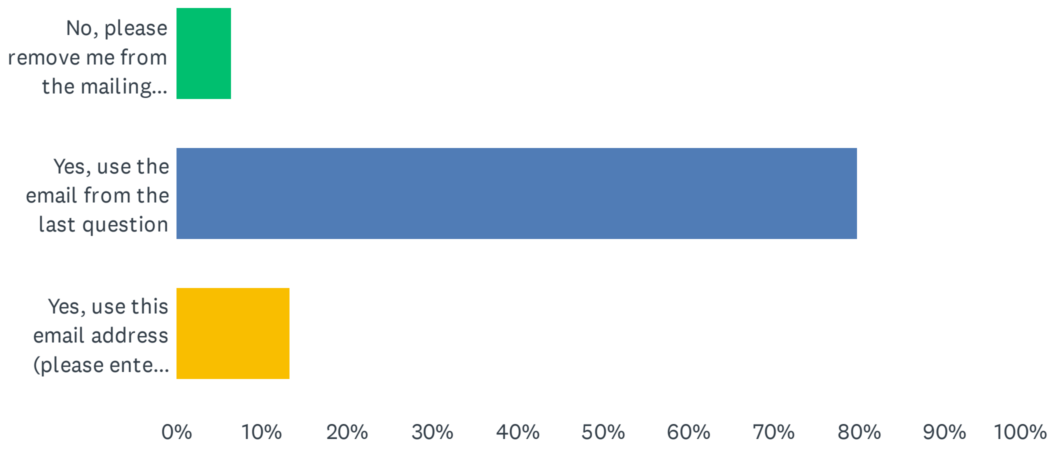


| ANSWER CHOICES | RESPONSES | |
|---|---|---|
| No, please remove me from the mailing list (please enter your email below, so we can remove it from the list) | 6.67% | 1 |
| Yes, use the email from the last question | 80.00% | 12 |
| Yes, use this email address (please enter below) | 13.33% | 2 |
| TOTAL | | 15 |

| # | EMAIL | DATE |
|---|---|---|
| 1 | [redacted] | 6/11/2024 1:30 PM |
| 2 | [redacted] | 6/11/2024 12:38 AM |
| 3 | [redacted] | 6/6/2024 1:48 PM |
